# Loss of a plant receptor kinase recruits beneficial rhizosphere-associated *Pseudomonas*

**DOI:** 10.1101/2020.11.02.364109

**Authors:** Yi Song, Andrew J. Wilson, Xue-Cheng Zhang, David Thoms, Reza Sohrabi, Siyu Song, Quentin Geissmann, Yang Liu, Lauren Walgren, Sheng Yang He, Cara H. Haney

## Abstract

Maintaining microbiome structure is critical for the health of both plants^1^ and animals^2^. In plants, enrichment of beneficial bacteria is associated with advantageous outcomes including protection from biotic and abiotic stress^3,4^. However, the genetic and molecular mechanisms by which plants enrich for specific beneficial microbes without general dysbiosis have remained elusive. Here we show that through regulation of NADPH oxidase, *FERONIA* kinase negatively regulates beneficial *Pseudomonas fluorescens* in the *Arabidopsis* rhizosphere microbiome. By rescreening a collection of *Arabidopsis* mutants that affect root immunity under gnotobiotic conditions, followed by microbiome sequencing in natural soil, we identified a *FERONIA* mutant (*fer-8*) with a rhizosphere microbiome enriched in *P. fluorescens* without phylum-level dysbiosis. Using microbiome transplant experiments, we found that the *fer-8* microbiome was beneficial and promoted plant growth. The effect of *FER* on rhizosphere Pseudomonads was independent of its immune coreceptor function, role in development, and jasmonic acid autoimmunity. We found that the *fer-8* mutant has reduced basal levels of reactive oxygen species (ROS) in roots and that mutants deficient in NADPH oxidase showed elevated rhizosphere Pseudomonad levels. Overexpression of the *ROP2* gene (encoding a client of FER and positive regulator of NADPH oxidase^5^) in *fer-8* plants suppressed Pseudomonad overgrowth. This work shows that *FER*-mediated ROS production regulates levels of beneficial Pseudomonads in the rhizosphere microbiome.

---

Eukaryotes are associated with communities of symbiotic microorganisms (the microbiome) that affect host heath and fitness^6^. In plants, the root-associated (rhizosphere) microbiome affects plant growth^7^, nutrient acquisition^8^ and resistance to both biotic and abiotic stresses^9–11^. While dysbiotic microbiomes have been associated with disease in both plants^1^ and animals^2^, in plants, enrichment of specific microbial taxa has been associated with growth promotion and pathogen protection. For instance, disease outbreaks can induce the assembly of beneficial microbes in the rhizosphere to enhance resistance to future disease^12,13^. Similarly, artificial enrichment of beneficial taxa in the lab or in the field can promote growth and protect plants from biotic and abiotic stresses^4,7,14^. These observations suggest that plants may be able to regulate the abundance of a few beneficial taxa to maximize fitness. However, the genetic regulators and mechanisms used by plants to control the abundance of beneficial microbes are largely unknown.

*Pseudomonas fluorescens* includes well-studied plant growth-promoting (PGP) and biocontrol strains^15,16^. Interestingly, *P. fluorescens* and related species are routinely enriched in agricultural disease suppressive soils. For example, disease suppressive soils that confer resistance to *Rhizoctonia solani*^13^, wheat take all disease^17^, *Fusarium* wilt^18^ and black root rot^19^ have all been associated with the enrichment of fluorescent Pseudomonads in the soil. The recurring observation that enrichment of Pseudomonads is associated with pathogen protection suggests that plants may possess a mechanism to specifically recruit beneficial *Pseudomonas* spp.^20–22^. In addition, certain natural accessions of *Arabidopsis thaliana* support different levels of rhizosphere *Pseudomonas* spp. while maintaining overall highly similar microbiome compositions^23^, indicating plants may have genetic mechanisms to regulate levels of beneficial *Pseudomonas* spp.

## *HSM13/FERONIA* inhibits rhizosphere *Pseudomonas* growth

To identify plant genes that regulate beneficial *Pseudomonas* spp. levels in the rhizosphere, We made use of 16 *hsm* (*<u>h</u>ormone mediated <u>s</u>uppression of <u>M</u>AMP triggered immunity*) mutants identified from a previous genetic screen^24^. The *hsm* mutations affect root immunity, and thus provide a genetic toolkit to identify novel genes that shape the rhizosphere microbiome. We screened these 16 *hsm* mutants^24^ for their ability to support growth of the beneficial *P. fluorescens* strain WCS365 expressing the luciferase operon (*P. fluorescens* WCS365-Luc) using a 48-well plate gnotobiotic system (Extended Data Fig. 1)^23^. We found that the *hsm13* mutant harbored consistently higher levels of rhizosphere *P. fluorescens* WCS365 relative to the parental line (Extended Data Fig. 2).

To test whether increased levels of *P. fluorescens* in the *hsm13* rhizosphere also occur in the presence of a complex microbial community, we grew *hsm13* in natural soil (Methods, Fig. 1a), and plated rhizosphere samples (normalized to the rhizosphere weight) on King’s B media to quantify fluorescent Pseudomonads^25^ (Fig. 1b). We found that the fluorescent Pseudomonads per gram of rhizosphere were enriched more than 10-fold in the rhizosphere of *hsm13* relative to wildtype plants (Fig. 1, b and c). We found that *hsm13* is stunted in both natural soil and axenic plates (Fig. 1a, Extended Data Fig. 3, a and b), and has a root hair elongation defect (Extended Data Fig. 3, c-e). To test whether plant morphological changes affect rhizosphere Pseudomonad levels, we tested *bri1-5* (a stunted mutant unable to perceive the growth hormone brassinolide^26^; Fig. 1a), and *ark1-1* (a root hair mutant with altered microtubule dynamics^27^). We found that both the *bri1-5* and *ark1-1* mutants have similar levels of rhizosphere fluorescent Pseudomonads as wildtype plants (Fig. 1c), suggesting that developmental defects are unlikely to underlie the enrichment of fluorescent Pseudomonads in *hsm13*.

**Fig. 1.**
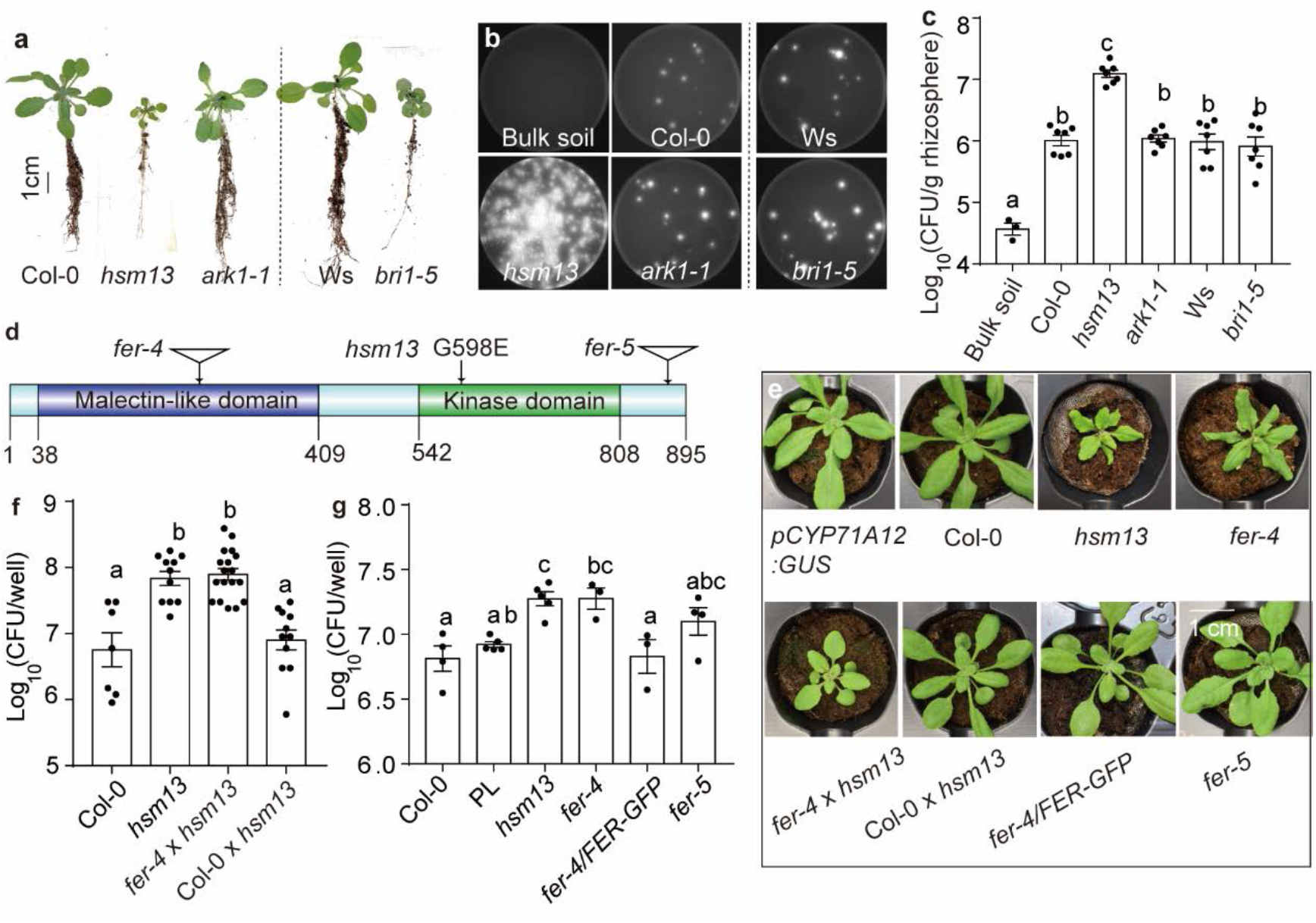
*hsm13/fer-8* harbors a high level of rhizosphere Pseudomonads due to a missense mutation in the *FERONIA* receptor kinase. (**a**) Morphology of wildtype plants (Col-0 and Ws ecotypes), and mutants *hsm13, ark1-1* (Col-0 background), and *bri1-5* (Ws background) when grown in natural soil; (**b**) *hsm13* harbored a high level of root-associated fluorescent Pseudomonads when grown in natural soil, while other mutants with similar developmental defects do not affect rhizosphere fluorescent Pseudomonads levels. Rhizosphere samples were plated on King’s B and imaged under UV light. **(c)** Quantification of fluorescent colonies per gram of rhizosphere samples. n=3-7. **(d)** The FER protein domains and the insertion/mutation positions of alleles described in this study. **(e)** Phenotypes of 3-week-old wildtype plants (Col-0), parental line (PL; *pCYP71A12:GUS), hsm13, fer-4, fer-5*, F1 crosses (*fer-4×hsm13* or Col-0×*hsm13*) and *fer-4/FER-GFP*. **(f)** F1 crosses between *fer-4×hsm13* have high level of rhizosphere Pseudomonads relative to Col-0, while Col-0×*hsm13* F1s restored Pseudomonads levels similar to wildtype plants. n=7, 11, 17 and 11 from left to right. **(g)** *fer-4 and fer-5* mutants have elevated levels of Pseudomonads in a hydroponic seedling assay. Each point represents the average of >6 plants from a single experiment. ANOVA and Turkey’s HSD were used for **c**, **f**, and (**g**) to determine significance; different letters indicate p < 0.05. Mean ± SEM is shown.

To identify the mutation that results in enriched *Pseudomonas* spp. in *hsm13*, we crossed *hsm13* to Col-0 and performed sequencing-assisted mapping by bulk segregant analysis (Methods). We identified a genomic region with a high frequency of SNPs present in the *hsm13*-like segregant population on the short-arm of chromosome 3 (Extended Data Fig. 4). We identified four non-synonymous SNPs in that region including a G1793A missense mutation in the coding sequence of *AT3G51550* (*FERONIA, FER*) in *hsm13* resulting in a predicted G598E amino acid substitution in the kinase domain (Fig. 1d). The previously described *fer-4* mutant shows a similar stunted morphology (Fig. 1e) and root hair developmental defects^28^. The F1s of a *fer-4×hsm13* cross had high *P. fluorescens* levels and small plant size similar to *hsm13* indicating that *fer-4* is allelic to *hsm13* (Fig. 1f, Extended Data Fig. 4). The F1 progeny of a Col-0×*hsm13* cross exhibited the *P. fluorescens* levels and plant size of Col-0 plants, confirming that *hsm13* is recessive (Fig. 1f, Extended Data Fig. 4). Using the gnotobiotic system, we found that *fer-4* (CS69044) had a similarly high level of WS365-Luc as the *hsm13* mutant and that *fer-5* (Salk_029056C) had a 1.95-fold increase in WCS365-Luc (Fig. 1f), which is consistent with previous data suggesting that *fer-5* is a partial loss-of-function allele^29^. Expression of *FER* under its native promoter (p*FER:FER-GFP^5^*) in the *fer-4* mutant completely restored the normal plant morphology and rhizosphere *P. fluorescens* WCS365 growth to wildtype levels (Fig. 1e and g). Collectively our data indicate that *hsm13* (*fer-8* hereafter) carries a loss of function mutation in *FER* resulting in stunting and rhizosphere Pseudomonad overgrowth.

## Microbiome profiling of *fer-8*

To determine the effect of the *fer-8* mutation on the overall rhizosphere microbiome composition, we grew *fer-8* and wildtype plants in the presence of natural soil microbiota [soil was from the same site over two consecutive years, Experiment 1 (Fig. 2) and 2 (Extended Data Fig. 5)] and performed 16S rRNA-based microbiome profiling. Samples from different years were clearly separated in the pooled PCoA analysis (Extended Data Fig. 5a) suggesting that the starting soil has the highest effect on rhizosphere microbiome composition. Despite the differences in soil composition from year-to-year, a consistently distinct microbiome composition was observed in *fer-8* relative to wildtype plants revealed by unconstrained principal coordinate analysis (PCoA), and 13.9%-18.2% of the differences in samples could be explained by plant genotype (PC2) (Fig. 2a; Extended Data Fig 5b). We observed no consistent shift in phylum level relative abundance between *fer-8* and wildtype plants (Fig. 2b; Extended Data Fig. 5). The *fer-8* microbiome had lower richness (number of Operational taxonomic units, OTUs) and Shannon diversity (a metric of species richness and evenness) compared to wildtype plants (Fig. 2c; Extended Data Fig. 5c)^30^. Although several bacterial families were enriched and depleted in each experiment, only Pseudomonadaceae were enriched in both experiments (Fig. 2d; Extended Data Fig. 5e). These data indicate that the Pseudomonadaceae are robustly enriched in the *fer-8* rhizosphere microbiome without phylum-level dysbiosis.

**Fig. 2.**
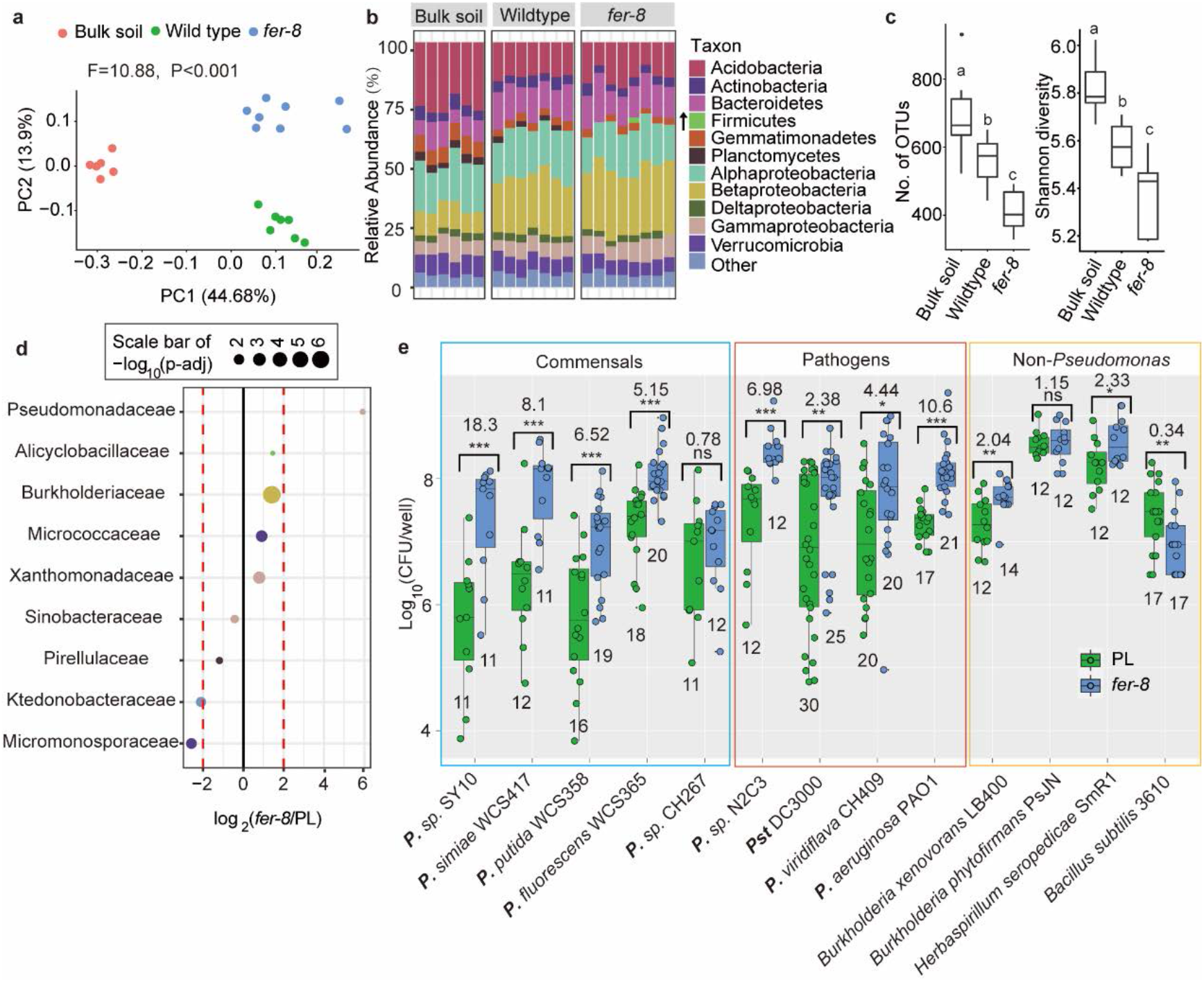
Pseudomonadaceae are enriched in the rhizosphere microbiome of *fer-8*. (**a)** Principal coordinate analysis (PCoA, based on the relative abundance of OTUs) of bulk soil and rhizosphere samples of *fer-8* and wildtype (Col-0) from Experiment 1. n=6-8. **(b)** Relative abundance of bacterial phyla or classes (for Proteobacteria) in bulk soil and rhizosphere samples. The arrow shows that only Firmicutes were significantly enriched in the rhizosphere of *fer-8* relative to the parental line (PL; 3.53-fold, calculated by DESEQ2). **(c)** Number of OTUs and Shannon diversity indexes in the bulk soil and rhizosphere samples. ANOVA and Turkey’s HSD were used to determine statistical significance, p < 0.05. **(d)** Significantly differentially abundant families between *fer-8* and wildtype. Colors show the taxonomic information for each family, and the dot size indicates the −log_10_ transformed adjusted p value for taxa with p<0.1. **(e)** Quantification of individual bacterial strains grown in the rhizosphere of *fer-8* and the PL under gnotobiotic conditions. Student’s t test was used to compare the significance between *fer-8* and PL for each strain (*p<0.05, **p<0.01, ***p<0.001). Boxplots (with median and first and third quartiles) and each data point are shown. Numbers denote the number of biological replicates over 2-3 independent experiments.

The genus *Pseudomonas* includes both beneficial microbes and plant pathogens^31^. This raises the question of whether *fer-8* specifically enriches beneficial *Pseudomonas* spp., or whether pathogenic *Pseudomonas* spp. are also enriched in the rhizosphere. We selected several phylogenetically diverse *Pseudomonas* strains (including both opportunistic pathogens and beneficial microbes), along with distantly related bacterial isolates, and tested whether they are enriched in the *fer-8* rhizosphere^31^. Commensal *Pseudomonas* strains included *P. putida* WCS358, *Pseudomonas* sp. CH267, *P. simiae* WCS417, *P. fluorescens* WCS365 and *Pseudomonas* sp. SY10 (identified from the natural soil used in this study). Pathogenic or opportunistic pathogens included *P. syrinage* pv. tomato (*Pst*) DC3000, *Pseudomonas* sp. N2C3^31^, *P. viridiflava* CH409^32^ and *P. aeruginosa* PAO1. We found that 8 of the 9 tested *Pseudomonas* strains (with the exception of commensal strain *Pseudomonas* sp. CH267) were enriched between 2- and 18-fold in the rhizosphere of *fer-8* relative to wildtype plants (Fig. 2e). All *non-Pseudomonas* strains tested, including *Burkholderia xenovorans* LB400, *B. phytofirmans* PsJN, *Herbaspirillum seropedicae* SmR1 and *Bacillus subtilis* 3610, exhibited 2-fold or lower enrichment, or were depleted, in the rhizosphere of *fer-8* (Fig. 2e). Collectively, these data suggest that the *fer-8* mutation robustly enriches most *Pseudomonas* spp.

## The *fer-8* microbiome is beneficial

Enrichment of fluorescent Pseudomonads in the rhizosphere is reminiscent of disease suppressive and growth promoting soils^13,15,33^. To test whether the *fer-8* associated microbiome is beneficial, we performed microbiome transplant experiments. We grew *fer-8* and its parental line in natural soil for 4 weeks (1^st^ generation, 2 plants per pot, Fig. 3a). The soil from *fer-8* or the parental line was then re-planted with wildtype plants. In the rhizosphere samples from the 1^st^ generation plants, we found an enrichment of fluorescent Pseudomonads in *fer-8* relative to its parental line (Fig. 3b and d). We observed a significant growth promotion effect of 2^nd^ generation plants grown in the presence of a *fer-8* microbiome (Fig. 3 a, c and e, Extended Data Fig. 6). Since the beneficial effects by *Pseudomonas* spp. have been shown to be dose-dependent^34^, we examined whether the growth promotion effect in the 2^nd^ generation plants correlated with the abundance of fluorescent Pseudomonads in soil. We found a significant positive correlation between the abundance of fluorescent Pseudomonads in the rhizosphere of 1^st^ generation plants (the soil used for 2^nd^ generation growth) and the biomass (shoot weight) of 2^nd^ generation plants (Pearson’s correlation, r = 0.97, p= 0.034, Fig. 3f). These data suggest that a single mutation in *FER* shifts the soil microbiome into one that promotes growth for the next generation of plants.

**Fig. 3.**
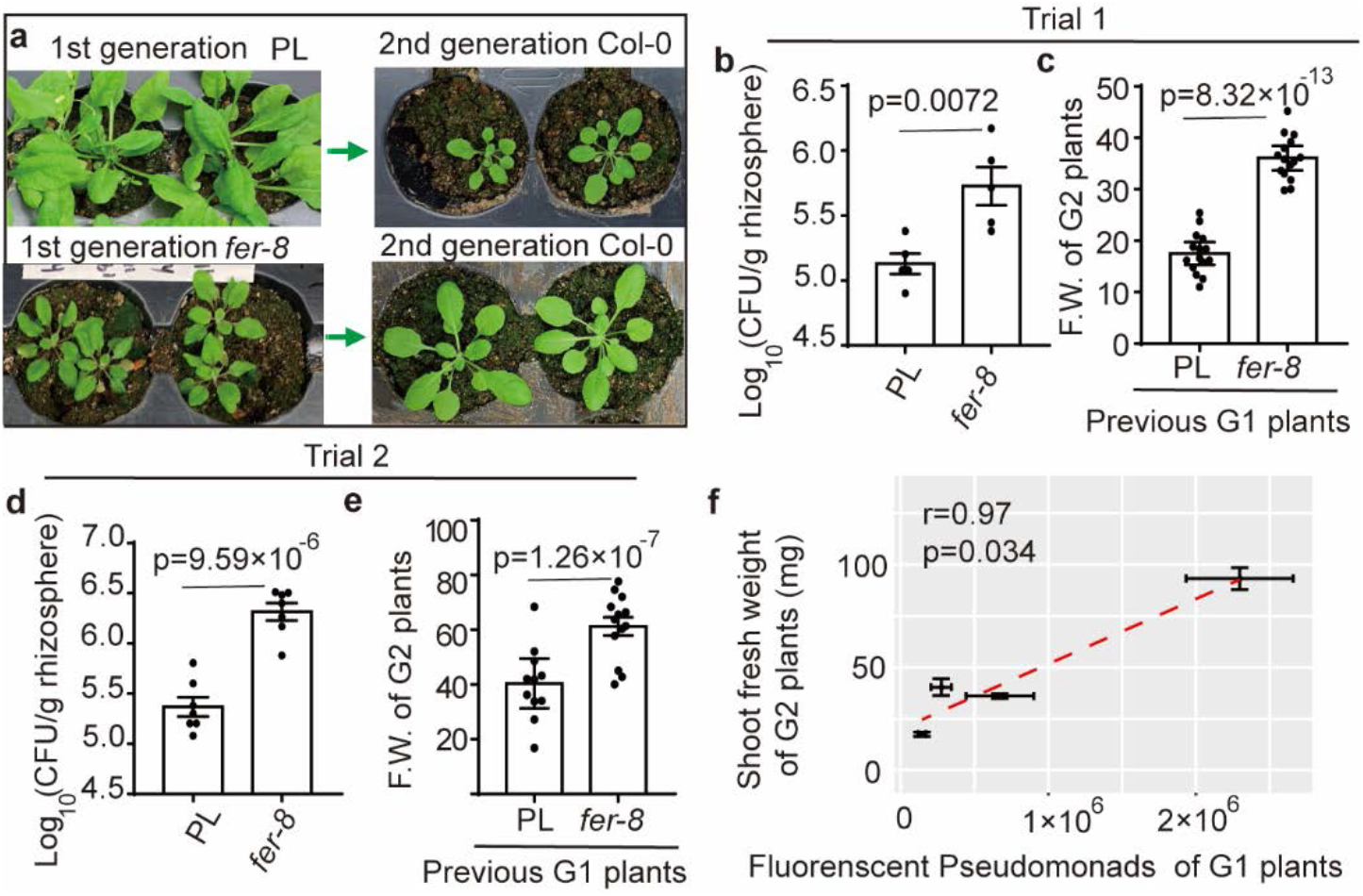
The *fer-8* rhizosphere microbiome confers growth promotion to the next generation of plants. **(a)** Representative images of the parental line (PL) and *fer-8* grown in natural soil (generation 1, G1) and wildtype Col-0 plants (generation 2, G2) are shown. **(b)** Fluorescent Pseudomonads were quantified in the rhizosphere of the parental line (PL) and *fer-8* in G1 plants after 4 weeks of growth, n=5. **(c)** The shoot fresh weight (F.W.) of G2 wildtype plants grown in the soil from microbiome transplants from the PL- or *fer-8-cultivated* soil, n=15. **(d-e)** A second replicate of the experiments shown in (**b-c)** was performed, n=7 for (**d**), n=11 and 13 for PL and *fer-8* in (**e**). Student’s t tests were used to determine statistical significance in (**b-e**). **(f)** The average fluorescent CFU counts from the first generation PL and *fer-8* plants were plotted against the average shoot F.W. of the next generation of plants grown in the same soil. A linear trend line (red dashed line) and Pearson’s correlation r are shown. Each data point represents the average value of all plants from one independent experiment from (**b**) to (**e**) with ±SEM.

In agriculture, suppressive soils are associated with enrichment of phylogenetically diverse *Pseudomonas* spp. While plants might have mechanisms to specifically enrich for beneficial strains, another possibility is that in the presence of the rhizosphere microbiome, that enrichment of pathogenic *Pseudomonas* may not be harmful due to competition with commensals in soil. *Pseudomonas* spp. are primarily associated with diseases of above-ground plant tissues from bacterial leaf spot to pith necrosis. However, we found that pathogenic strains *Pst* DC3000 and *Pseudomonas* sp. N2C3 robustly cause stunting when added to the roots of gnotobiotic plants (Extended Data Fig. 7 a and b). To test if these strains can cause disease in soil, pathogenic *Pst* DC3000 and *Pseudomonas* sp. N2C3 were inoculated in the rhizosphere of plants grown in natural soil. Neither strain caused disease symptoms after inoculation to a final concentration of 10^5^ and 10^6^ CFU/gram of soil (Extended Data Fig. 7 c and d). These data suggest that the enrichment of *Pseudomonas* pathogens may not efficiently cause disease in the presence of a natural soil community so general enrichment of rhizosphere *Pseudomonas* might not present a risk of disease.

## Jasmonic Acid (JA) signaling and innate immune receptors do not affect *Pseudomonas* enrichment

To reveal transcriptional changes in the *fer-8* mutant that could explain the increase in *Pseudomonas* colonization, we performed transcriptional profiling in both shoots and roots from *fer-8* and the parental line (Supplementary Table 1). We identified 675 up-regulated genes in the shoots of *fer-8* relative to wildtype plants (Supplementary Table 2). Surprisingly, we found only 82 up-regulated genes in *fer-8* roots relative to the parental line, and there were no significantly enriched GO terms. In contrast, we found that the genes upregulated in shoots were enriched in GO terms related to defense, response to fungi, and JA signaling (Extended Data Fig. 8, Supplementary Table 3), consistent with previous reports of JA activation in the shoots of *fer-4*^28^. *fer-8* exhibited enhanced resistance to the fungal pathogen *Botrytis cinerea*, shoot-specific expression of JA responsive genes, and enhanced anthocyanin accumulation in petioles (Extended Data Fig. 8). The transcriptional changes in shoots were largely limited to JA signaling while expression of other hormone signaling pathways were relatively similar between *fer-8* and the parental line (Extended Data Fig. 9, Supplementary Table 4). These data suggest that the *fer-8* mutation results in activation of JA signaling in shoots but not in roots.

While we did not observe changes in JA signaling in the root transcriptome, we hypothesized that a shoot-to-root JA-dependent signal could affect the rhizosphere microbiome. To test whether JA-mediated auto-immunity in shoots affects rhizosphere *Pseudomonas* colonization in *fer-8*, we constructed a double mutant with *coi1-16* (deficient in JA perception^35^) and *fer-8* (*coi1-16 fer-8*). We found that the *coi1-16 fer-8* mutant suppressed the stunting phenotype of *fer-8* (Extended Data Fig. 10). However, the *coi1-16 fer-8* double mutant retained enhanced *Pseudomonas* growth similar to the *fer-8* single mutant (Fig. 4a). Enhanced JA in shoots might antagonize salicylic acid (SA) signaling^36^. To determine if effects on SA signaling could explain the enhanced levels of *P. fluorescens* in the *fer-8* mutant, we tested rhizosphere *Pseudomonas* levels in the SA perception deficient mutant (*npr1-1*)^37^, the SA biosynthesis mutant (*sid2-1*)^38^ and the SA auto-immune mutant (*snc1*)^39^. We found no significant changes in rhizosphere fluorescent Pseudomonads in SA mutants (Extended Data Fig. 11). These data collectively indicate that neither JA auto-immunity nor SA-JA antagonism fully explains the increase in rhizosphere *Pseudomonas* colonization in the *fer-8* mutant.

**Fig. 4.**
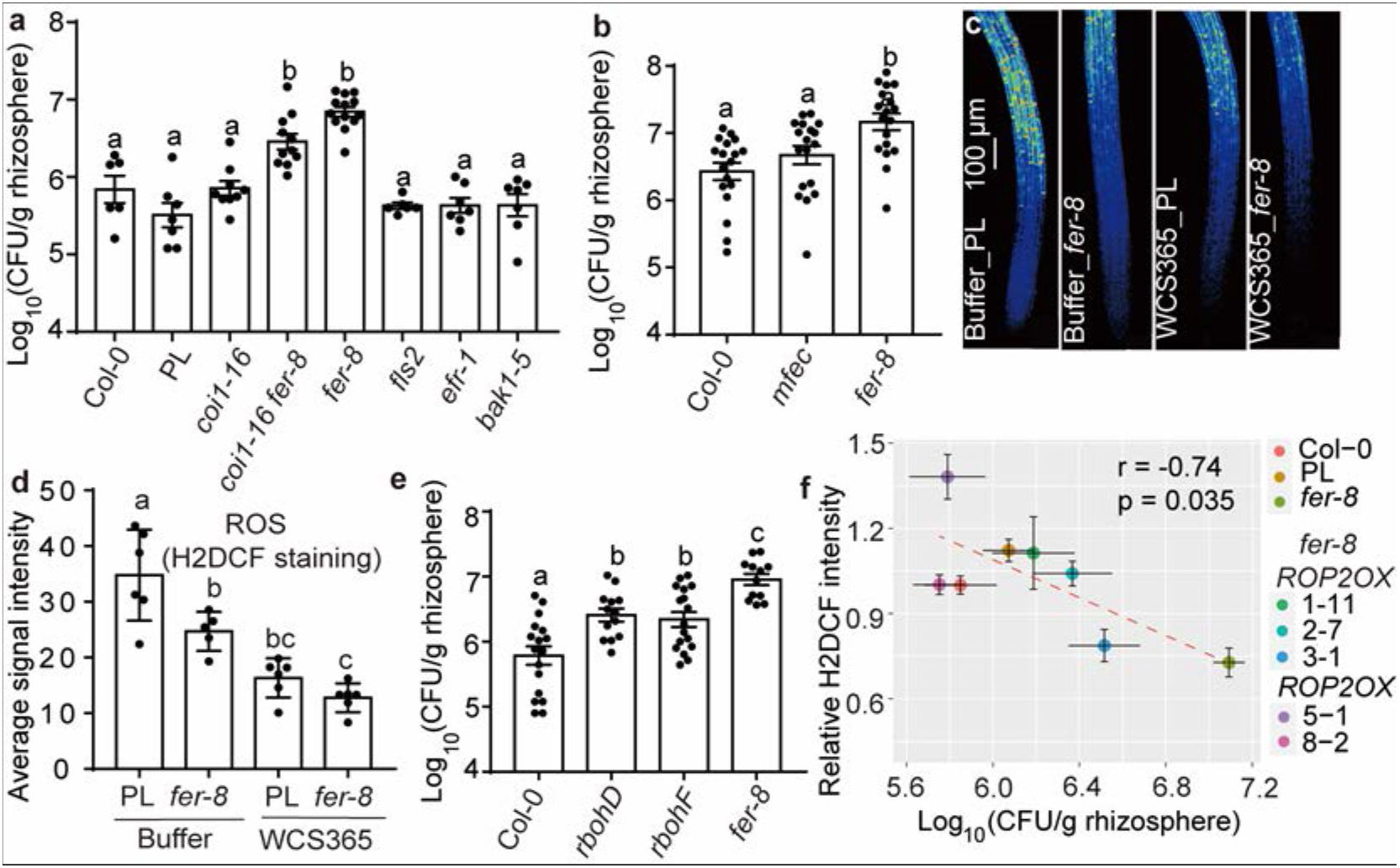
Root ROS negatively regulates rhizosphere *Pseudomonas*. **(a)** Mutants deficient in immune receptors that are interaction partners of FER (*fls2, efr-1, bak1-5*), and the JA receptor (*coi1-16*) do not affect rhizosphere *Pseudomonas* levels, and *fer-8 coi1-16* double mutant does not restore the *fer-8 Pseudomonas* overgrowth phenotype. n = 6, 7, 9, 11, 13, 6, 7, 7 from left to right. (**b**) A quadruple mutant deficient in *min7, fls2, efr* and *cerk1* (*mfec*) does not change rhizosphere Pseudomonad level as *fer-8*. n=18 from three independent experiments. (**c**) Representative images of H2DCF-DA (ROS-binding dye) stained roots of the parental line (PL) and *fer-8* pre-treated with buffer or *P. fluorescens* WCS365. (**d**) Quantified average H2DCF-DA signal intensity in roots from two independent experiments. (**e**) Mutants deficient in NADPH oxidase (*rbohD/F*) show elevated rhizosphere fluorescent Pseudomonads when grown in soil. n=17, 13, 17 and 12 from left to right (3-4 independent experiments). (**f**) The average log_10_(fluorescent CFU/g rhizosphere) from different genotypes were plotted against the average relative H2DCF staining signaling intensity. A linear trend line (red dashed line) is shown and Pearson’s correlation was used to determine significance (p=0.035). Different letters indicate p< 0.05 by ANOVA and Tukey’s HSD in a, b, d and e.

FER interacts with immune receptors and facilitates the complex formation of innate immune receptor complexes that include EF-TU RECEPTOR (EFR) and FLAGELLIN-SENSING 2 (FLS2) with their co-receptor BRASSINOSTEROID INSENSITIVE 1–ASSOCIATED KINASE 1 (BAK1)^40^. We reasoned that if FER regulates Pseudomonads through its immune scaffold function, then immune receptor mutants should also increase rhizosphere Pseudomonads. However, we found that *fls2, efr-1* and *bak1-5* mutants all have similar levels of rhizosphere fluorescent Pseudomonads as wildtype plants when grown in natural soil (Fig. 4a). A recent study found that a mutant deficient in multiple immune receptors and vesicle-trafficking (*mfec: min7 fls2 efr cerk1*) causes dysbiosis in the endophytic phyllosphere microbiome and decreased alpha diversity^1^. We wondered whether the *mfec* mutant would show rhizosphere enrichment of *Pseudomonas* similar to *fer-8*. Although the enrichment of Pseudomonads in the *fer-8* was reproducible in a distinct natural soil (from Michigan State, USA), we did not observe significant enrichment of Pseudomonads in the rhizosphere of *mfec* mutants (Fig. 4b). This indicates that the mechanism of rhizosphere microbiome changes in *fer-8* is distinct from the *mfec* mutant, and that innate immune perception as well as immune scaffold functions of FER are largely dispensable for the selective enrichment of Pseudomonads in *fer-8*.

## FER maintains root reactive oxygen species (ROS) to control Pseudomonads

In addition to the MAMP-triggered ROS burst (inducible ROS)^40^, FER also positively regulates basal ROS levels in roots through ROP2 (a small GTPase). ROP2 is a positive regulator of plasma membrane NADPH oxidases and *fer-4* and *fer-5* mutants have reduced basal ROS levels in roots^5^. We hypothesized that loss of FER might result in decreased basal ROS resulting in *Pseudomonas* overgrowth. By staining roots with the ROS-sensitive dye H2DCF-DA^5^, we found a significant decrease in basal ROS levels in buffer-treated *fer-8* compared to the parental line (Fig. 4c and d). Importantly, roots treated with *P. fluorescens* WCS365 exhibited a significant reduction in root ROS levels in both *fer-8* and the parental line (Fig. 4c and d), indicating that suppression of basal root-surface ROS might be crucial for *Pseudomonas* fitness in the rhizosphere.

To test if altering rhizosphere ROS levels could affect *Pseudomonas* growth in the rhizosphere, we tested *<u>R</u>espiratory <u>B</u>urst <u>O</u>xidase <u>H</u>omologues* mutants (*rbohD, rbohF*), which are deficient in the membrane-associated NADPH oxidase and dampen apoplastic ROS production^41^. We found that both *rbohD* and *rbohF* have significantly elevated rhizosphere fluorescent Pseudomonads in natural soil (Fig. 4e). To test if *FER* acts through NADPH oxidase to regulate rhizosphere *Pseudomonas*, we overexpressed the FER interacting client *ROP2* in *fer-8*, which has previously been shown to enhance ROS in a *fer-5* mutant^5^. We found that *fer-8 ROP2OX* significantly increased root ROS levels relative to *fer-8* (H2DCF signal intensity) and decreased the level of fluorescent Pseudomonads observed in the *fer-8* rhizosphere (Extended Data Fig. 12). By plotting the average rhizosphere fluorescent Pseudomonads levels against the average ROS levels (measured by H2DCF signal intensity) in different genetic backgrounds, we saw a significant negative correlation (r = −0.74; p = 0.035) between ROS and Pseudomonad levels (Fig. 4f). Collectively these data indicate that *ROP2*-mediated regulation of ROS is required for *FER* mediated regulation of rhizosphere Pseudomonads.

## Discussion

This work revealed that *FER* regulates rhizosphere *Pseudomonas* colonization through suppressing NADPH oxidase activity. We found that genetic manipulation of ROS production in plants can shape the microbiome and gate access of beneficial commensals to the rhizosphere. Although the ROS burst is a critical step of innate immune activation in plants, we found that FER-mediated regulation of rhizosphere Pseudomonads is dependent on basal ROS rather than inducible ROS triggered by MAMP perception and innate immune receptors. From an evolutionary standpoint, this could be because roots are constitutively exposed to a MAMP-rich environment and are less sensitive to MAMPs^42^. Moreover, rhizosphere microbes can suppress pattern-triggered immunity (PTI)^43^ and thus plants may rely on basal ROS levels to gate rhizosphere *Pseudomonas* colonization.

It is unclear why decreased ROS in *hsm13* relatively specifically enriches *Pseudomonas* spp. as ROS is toxic to most microbes. We speculate that root surface ROS only affects a localized region close to the root surface (rhizoplane), which may be a region where beneficial *Pseudomonas* spp. are specifically enriched^23^ and therefore may be more vulnerable to plant defenses than other taxa. Spatial-temporal resolution of rhizosphere communities may reveal why manipulation of root ROS effectively gates colonization by *Pseudomonas* spp.

The enrichment of *Pseudomonas* is of agricultural interest because it is frequently associated with disease suppressive and growth promoting soils. *FER* is a versatile receptor kinase that positively regulates growth and immunity, and is thus a potential target that could be hijacked by pathogens. Both the fungal pathogen *Fusarium oxysporum* and nematodes secrete functional RALF-like peptides to manipulate FER activity and to promote pathogenesis^44,45^. Interestingly, root associated *Pseudomonas* spp. are enriched in disease suppressive soil towards either *Fusarium oxysporum* or nematodes^18,46^ suggesting that pathogen manipulation of *FER* signaling in crops^47,48^ is a possible mechanism for increased *Pseudomonas* after pathogen attack. Collectively this work has the potential to guide novel breeding and microbiome engineering practices in agriculture through manipulation of FER signaling.

## Methods

### Plant materials and growth methods

*Arabidopsis* seeds were surface sterilized (washed in 70% ethanol for 2 min, 10% bleach for 2 min, and 3 times in sterile water), and stored at 4°C for at least 2 days before use. For assays on plates with solid media, seedlings were grown on 1/2× MS medium with 1% phytoagar and 1% sucrose. Plates were grown at 22°C under 90-100 μE light on a 12 hours light / 12 hours dark cycle. The *Arabidopsis* Col-0 ecotype was used as the wildtype genetic background in this work, *fis2*^49^, *efr-1*^50^, *bak1-5*^51^, and *min7fls2 efr cerk1* (*mfec*)^1^ were reported previously. The *ark1-1* mutant has a 35Spro:*EB1b*-GFP reporter, which does not alter the root hair phenotype^27^.

### Gnotobiotic rhizosphere bacterial quantification assay

The assay (Extended Data Fig. 1) was performed as described previously^23^. Briefly, seeds were germinated on Teflon mesh disks in 300 μL MS media with 2% sucrose in 48-well plates. After 10 days, the media was changed to 270 μL media without sucrose (1×MS, 0.1% MES buffer) so that bacteria rely on plant root exudate as a carbon source. After 2 more days, 30 μL of *P. fluorescens* WCS365 transformed with the lux operon from *Aliivibrio fisheri* (WCS365-Luc hereafter) was added. Two days after inoculation, media containing bacteria from the rhizosphere in 48-well plates was transferred to opaque white 96-well plates prior to reading to avoid background from the plants and Teflon mesh, and Luc photo counts were measured using a Spectra Max i3 plate reader (Molecular Devices). Any plants with translucent or water-soaked leaves were discarded from the assay and not used for bacterial treatment. To generate WCS365-Luc, a transposon containing the *Aliivibrio fisheri* LUX operon was integrated into the *P. fluorescens* WCS365 genome by conjugation with *Escherichia coli SM10pir* containing pUT-EM7-LUX^52^. To ensure that the insertion did not affect WCS365 growth promotion ability, we confirmed that WCS365-Luc promoted lateral root growth to a similar level as wildtype WCS365 (Extended Data Fig. 1). We found a linear relationship between the WCS365-Luc bacterial CFU counts and the Luciferase signal (Extended Data Fig. 1), indicating that the WCS365-Luc strain can be used to approximate bacterial numbers in the rhizosphere.

### Mapping of *hsm13*

The (*FER*) gene was cloned by bulk segregant analysis^53^. We hypothesized that the same mutation might cause both stunting and rhizosphere *Pseudomonas* enrichment in *hsm13*, and so we used stunting to screen for *hsm13-like* segregants. Briefly, we backcrossed *hsm13* to wildtype plants (Col-0) and identified 30 stunted F2 segregates (*hsm13* like) from 140 F2 plants. We found that all F3s from these lines were stunted indicating they were homozygous for the mutation leading to stunting. We then sampled 90 plants (3 plants from each F2 line, one leaf per plant) and extracted DNA from each leaf separately. Genomic DNA samples were quantified using Quant-iT^™^ PicoGreen^™^ dsDNA Reagent (Invitrogen). DNA from different samples was mixed at equimolar ratios. A 1:1 mix of DNA from Col-0 and *pCYP71A12:GUS* (the parental line of *hsm*13) was also sequenced as a reference sample. Paired-end sequencing (150 bp reads) was performed on an Illumina HiSeq by Novogene. After filtering, approximately 35 million single-end reads (approximately 37X coverage after trimming, mapping and filtering) and 34 million single-end reads (approximately 37X coverage) were mapped to the TAIR 10 *Arabidopsis* reference genome for the *hsm13* segregates population and the pooled reference samples, respectively^54^. After SNP calling relative to the TAIR10 reference genome, 333 and 486 non-synonymous SNPs were identified in *hsm13* segregates and the pooled reference samples, respectively. P_SNP_ was calculated as Mutant SNP / (Mutant SNP+ wildtype SNP) in Excel. SNPs present in both *hsm13* segregates and the pooled reference samples were discarded from further analysis.

### Harvesting rhizosphere samples

Natural soil for the majority of experiments (except Fig. 4b) was harvested from the UBC farm (49°15.0’N, 123°14.4’W), Vancouver, British Columbia, Canada. This is a disturbed site on the UBC farm that was naturally colonized by wild *Arabidopsis thaliana*. The top 10-20 cm of soil was collected and sieved (3-mm sieve) to remove rocks, insects and plant debris. *Arabidopsis* seedlings were grown on ½ x MS plates with 1% sucrose for 8-10 days before transplanting to soil. We blended additional inorganic growth materials and soils to improve drainage and plant health. The final soil substrate consisted of 1:0.5:1 natural soil: calcine clay (Turface): perlite for microbiome sequencing and other studies using natural soil. Both rhizosphere and bulk soil samples were harvested at 17-20 days after transplanting.

The experiment to quantify fluorescent *Pseudomonas* in the rhizosphere of *hsm13* and *mfec* (Fig. 4b) was performed in natural soil collected in Michigan, USA. Plants were grown in *“Arabidopsis* Mix” greenhouse potting soil [equal parts of Suremix (Michigan Grower Products), medium vermiculate and perlite] and was autoclaved once before use. Individual pots were supplemented with natural soil slurry prepared from a soil where wild accessions of *Arabidopsis* were found at Michigan State University’s Southwest Michigan Research and Extension Center (Benton Harbor). To prepare soil slurry, 25g of the soil was mixed with 1 liter of autoclaved ddH2O for 30 min on an orbital shaker, filtered through a 70-um cell strainer and 20 mL of soil slurry was supplemented to each pot uniformly by top irrigation. Plants were grown under relative humidity set at 50%, temperature at 22 °C, light intensity at 100 μE m^−2^ s^−1^ and photoperiod at a 12:12 hours dark: light cycle. Four-week-old plants were used for rhizosphere sampling based on the protocol below.

To collect rhizosphere samples, we collected roots and closely-adhered soil. To harvest rhizosphere samples, pots were inverted to transfer the soil and whole plant to a gloved hand. The soil was then gently loosened from the root until just the roots and closely-adhered soil remained. Gloves were cleaned with 70% ethanol between samples, and fresh gloves were used between genotypes. Rhizosphere samples were weighed and buffer was added to 0.05 g/mL (7.5 mM MgSO4 and 20% glycerol). Samples were homogenized using a Tissuelyser (2 x 90 seconds at 25 rps). Samples were serially diluted and 100 μl of the bulk soil (0.0025 g/ml) or rhizosphere samples (0.00025 g/mL) were plated on King’s B plates and imaged using a UV light source.

### Rhizosphere microbiome transplant assay

For microbiome transplant experiments, we grew first-generation plants (either *fer-8* or the parental line) for 3.5-4 weeks (2 plants/pot) in natural soil to allow assembly of a genotype-specific rhizosphere microbiome. We then cut the shoots of the first-generation plants and thoroughly mixed all soil from same genotype together in a sterilized container. We then immediately (the same day) put the mixed soil (with genotype-specific microbiomes) into new clean pots to grow second-generation plants. The trays and growth chamber were sterilized with 70% ethanol before the experiment and all plants were watered with autoclaved water for both the first and second generation plants. Different genotypes were put in separate trays side by side in the same growth chamber and grown under 80-100 μE light on a 12 hour light/12 hour dark cycle. For first generation plants, about 15% of the plants were bolting 4 weeks after transplanting, we only chose non-bolting plants for the rhizosphere sampling to avoid effects of differences in developmental stage. For both generation 1 and 2 plants, 1L of 1/4x Hoagland’s fertilizer was added to each tray.

### 16S rRNA microbiome sample preparation, sequencing and analysis

For microbiome sequencing, four individual rhizosphere or bulk soil samples were pooled as one replicate. Sample processing and sequencing was performed as described in the Earth Microbiome Project Illumina 16S rRNA protocol (www.earthmicrobiome.org). Briefly, total soil or rhizosphere DNA was extracted using the PowerSoil^®^ DNA Isolation kit (MoBio Laboratories, Carlsbad, CA). DNA concentrations were determined using a Quant-iT^™^ PicoGreen^™^ dsDNA Assay Kit. Paired end 300 bp sequencing was performed on an Illumina MiSeq. Adaptor sequences were trimmed with cutadapt and DADA2 was used to generate an amplicon sequence variant (ASV) table^55^. The Qiime2 implementation of vsearch was used to bin ASVs at 97% identity and the q2-feature-classifier was used to assign taxonomy using a naive Bayesian approach. Principle covariate analysis was performed using Bray-Curtis dissimilarity of relative abundances (OTU level) with the vegan package in R. Differentially abundant families were identified using the DESEQ2 package^56^.

### Transgenic plants

To overexpress *ROP2*, a gene-specific primer pair was used to amplify the coding sequence of *ROP2* (Forward: 5’-ATA<u>TCTAGA</u>ATGGCGTCAAGGTTTATAAAGT-3’ and reverse: 5’-ATA<u>CTGCAG</u>TCACAAGAACGCGCAACGGTTC-3’), with the restriction enzyme sites for *Xba*1 and *Pst*1 added to the forward and reverse primers, respectively. The PCR product was digested by *Xba1* and *Pst*1 enzymes and subcloned into a binary vector pCambia1300^57^. The sequence was confirmed by Sanger sequencing, and the plasmid was introduced into *Agrobacterium* GV3101 for floral dip transformation of *Arabidopsis* (Col-0 and *fer-8*). T1 and T2 transformants were selected/confirmed in 1/2× MS with 1% sucrose and 50 μg/mL hygromycin.

### Plant genotyping

Primers for *fer-4*’are listed here^29^: P1 5’-GATTACTCTCCAACAGAGAAAATCCT-3’, P2 5’-CGTATTGCTTTTCGATTTCCTA-3’, P3 5’-ACGGTCTCAACGCTACCAAC-3’, P4 5’-TTTCCCGCCTTCGGTTTA-3’; and primers for Salk_029056C: LP 5’-TGGTAGGATTCCGTTAAAATGC-3’, RP 5’-CAGAGTATTTCAGACGGCAGC-3’, LB 5’-ATTTTGCCGATTTCGGAAC-3’. For detection of *ROP2OX* in T1 lines, we used a pair of primer targeting the *35S* promoter and *ROP2* gene (5’-CTATCCTTCGCAAGACCCTTC-3’; 5’-GCAACGGTTCTTATTCTTTTTCT-3’), respectively.

For *fer-8 coi1-16* double mutant, F2 progeny of *fer-8* × *coi1-16* were selected on ½× MS agar plates supplemented with 20 μM MeJA, and seedlings insensitive to JA-mediated root growth inhibition were selected as *coi1-16* homozygous lines, and *fer-8* allele specific SNP detection primers were designed by a web tool (http://ausubellab.mgh.harvard.edu/). The primer (5’-ACATCGTCATCTTGTGTCCT TGATGGG-3’, 5’-GGGTTCAAGGCTGGACGAGCG-3’) can specifically amplify wildtype *FER* fragment but not *FER^G598E^* allele (10-500 ng template DNA, elongation temperature: 57 degree for 25 cycles). The selected double mutants were confirmed by sanger sequencing.

### Rhizosphere growth of non-tagged commensals

Rhizosphere commensals strains were grown in the rhizosphere of *fer-8* and the parental line (*pCYP71A12:GUS*). All bacterial strains were cultured in LB broth or solid LB media without antibiotics at 29°C. *Pseudomonas* sp. SY10 was isolated by plating a rhizosphere sample from natural soil on King’s B, selecting a fluorescent colony and streaking for single colonies. The identity as a *Pseudomonas* sp. was determined by amplifying the 16S rRNA with primers 8F (5’-AGAGTTTGATCCTGGCTCAG-3’) and 1392R (5’-ACGGGCGGTGTGTRC-3’), and sequencing with primer 8F. To quantify CFUs, *fer-8* or the parental line were grown as described above for the gnotobiotic system and bacteria were added to a final OD_600_ = 0.00002 in 300 μl. The media containing bacteria surrounding plant roots was serially diluted and plated to calculate CFUs. For most of the strains, rhizosphere samples were serially diluted and plated on LB plates 2 days after inoculation, while samples of *Burkholderia phytofirmans* PsJN and *Herbaspirillum seropedicae* SmR1 were plated at 4 days after inoculation. *Pst* DC3000 and *P. viridiflava* CH409 were plated at 50-72 hours after inoculation due to slow growth in the rhizosphere. Bacterial strains used in this study were previously described: *Pseudomonas* sp. CH267^23^, *P. fluorescens* WCS365^58^, *P. simiae* WCS417^59^, *P. aeruginosa* PAO1^60^, *P. putida* WCS358^58^, *Bacillus subtilis* NCIB 3610^61^, *Burkholderia xenovorans* LB400^62^, *Herbaspirillum seropedicae* SmR1^63^, *Burkholderia phytofirmans* PsJN^64^, *P. syringae* pv. tomato (*Pst*) DC3000^65^, *Pseudomonas* sp. N2C3^66^, *P. viridiflava* CH409^32^. *Pseudomonas* sp. SY10 was identified in the natural soil used in this study.

### qRT-PCR

RNA was extracted using an RNAeasy isolation kit (Qiagen). RNA samples were treated with Turbo DNA-free (Ambion) kit and single-stranded cDNA was generated from 1-2 μg RNA using Superscript III (Invitrogen) and Oligo dT primers following the manufacturer’s instructions. Quantitative RT-PCR was performed on a 7,500 Fast Real-Time PCR machine (Applied Biosystems) using SYBRTM Green Master Mix kit (ThermoFisher). Primers for qRT-PCR are listed here^32^: *PDF1.2*, 5’-CTTATCTTCGCTGCTCTTGTTC-3’ and 5’-TGGGAAGACATAGTTGCATGAT-3’; *VSP1:* 5’-CTCAAGCCAAACGGATCG-3’ and 5’-TTCCCAACGATGTTGTACCC-3; *VSP2*, 5’-TCAGTGACCGTTGGAAGTTGTG-3’ and 5’-GTTCGAACCATTAGGCTTCAATATG-3’.

### RNA Sequencing and data analysis

For RNA sequencing, plants were grown on ½× MS with 1% phytoagar and 1% sucrose. For both the parental line (*pCYP71A12:GUS*) and *hsm13*, samples were harvested at 11 days after germination. Shoots from 2 plants or roots from 5 plants were pooled for each sample. RNA was extracted using a Qiagen RNAeasy isolation kit. RNA samples with a concentration higher than 300 ng/uL and RIN number high than 8 were used for library preparation. Construction of libraries and sequencing were performed at the Michael Smith Genome Sciences Center (http://www.bcgsc.ca/). Paired end 75 bp RNA-sequencing was performed using an Illuminia Hi-seq 2500 platform. High quality reads were mapped to the Tair 10 genome using Bowtie2^67^, and transcript quantification was performed with RSEM^68^. Differential expression analysis was performed in R (https://www.R-project.org/.) with the DESeq2 package^56^. Differentially expressed genes were filtered by p_adj_ < 0.1. GO enrichment analysis was done by AgriGO^69^. The core JA responsive genes were those that were induced in response to JA treatment at all timepoints from 1 to 16 hours (a total of 12 sampling points)^70^, and all other hormone responsive genes were obtained from a previous publication^71^. The Heatmap.2 function from gplots package in R were used to generate heatmaps.

#### Infection assay

*Botrytis cinerea* infection assays were performed as described^72^. Leaves 7-9 from 4-week old plants were excised and placed adaxial side down onto 1% agar plates for infection. 6 μL of 5 ×10^5^ spores/ml was dropped onto the abaxial leaf surface. Lesion diameters were measure at 3 days after inoculation.

For *Pst* DC3000 and *Pseudomonas* sp. N2C3 inoculation in soil, six-day-old *Arabidopsis* seedlings were grown on ½× MS agar with 1% sucrose and transferred into natural soil [1:0.5:1 natural soil: calcine clay (Turface): perlite]. Inoculation was performed 1 or 2 days after transplanting. Overnight bacterial cultures were centrifuged and washed in 10 mM MgSO_4_ three times before being diluted to an OD_600_ of 0.5 and 0.05. One milliliter of bacterial inoculum at OD_600_=0.5 or 0.05 for each pot (approximately 70 gram soil mixture) was added to the soil taking care to not touch the seedlings to reach a final concentration of 3×10^5^ and 3×10^6^ CFU/gram soil, respectively. The same volume of 10 mM MgSO4 was used as a buffer control. Different groups were grown side by side in the same tray, but were watered separately in different trays to avoid cross contamination.

For *Pst* DC3000 and *Pseudomonas* sp. N2C3 inoculation on plates, seeds were germinated on 1/2× MS plates (without sucrose), grown for 6 days, and inoculated with 5 μL bacteria (OD_600_=0.05). The same volume of 10 mM MgSO4 was used as a buffer control. The plates were scanned 7 days after inoculation, and the plants were weighed on the same day.

### Detecting ROS in roots

ROS detection was performed using 2’,7’-dichlorodihydrofluorescein diacetate (H2DCF-DA) fluorescent dye. Plants were grown on ½× MS agar plates supplemented with 1% sucrose for 4 days. Roots were inoculated with 3 μL of buffer (10 mM MgSO_4_) or *P. fluorescens* WCS365 (OD_600_=0.01), and seedlings were imaged 24 hours after inoculation. H2DCF-DA was dissolved in DMSO (10 mg/mL) and then diluted to a 500 mM stock (10×) in 0.1 M PB buffer (pH 7.0) and stored at −20°C. Before use, H2DCF-DA aliquots were thawed in the dark, stored on ice, and diluted to a 1× working concentration in 2 mL ½× MS media with 0.1% MES [2-(N-Morpholino) ethanesulfonic acid sodium salt]. Whole seedlings were transferred to a 12-well plate with 2 mL staining solution per well and stained for 15 minutes at room temperature in the dark. Imaging chambers were constructed according to a JOVE protocol^73^. Imaging chambers were molded from Poly(dimethylsiloxane) gel. A 1.5% agar pad was placed into a chamber, and roots were rinsed in ½× MS and mounted onto the pad. Glass strips placed at both ends of the glass slide provided consistent coverslip spacing and root positioning directly against the coverslip, which allowed for consistent optical resolution and fluorescence signal during image acquisition. Confocal images were acquired with a 10×/0.40 NA objective on a Leica SP8 laser scanning confocal microscope using a white light laser. H2DCF-DA was excited with a 504 nm laser and a HyD detector was used to capture emission between 511 nm - 611 nm, with detection time-gated with 0.3 – 12 ns to reduce autofluorescence. To ensure that all H2DCF-DA emitted fluorescence above the background was captured, large 3D image stacks (depth of ~120-150 microns) were taken at 2.408 μm steps (total of 50-60 images per root). Images were converted to .tiff files using FIJI^74^, and root area was traced manually. The H2DCF-DA signal density was quantified based on 2D maximum intensity image projections in FIJI and the total intensity was divided by the root area.

For H2DCF signal detection in *fer-8* ROP2OX lines (Extended Data Fig. 12), four day old seedlings (grown on ½× MS agar supplemented with 1% sucrose) were stained in a H2DCF solution as described above. Seedlings were transferred onto new ½× MS agar plates for imaging. A Leica M205FA fluorescence stereo microscope equipped with a Leica PLAN APO 2.0× CORR objective was used for high throughput imaging of two to four seedlings per image. Images were acquired with a GFP filter set (excitation filter ET470/40 nm-emission filter ET525/50 nm) at 2 or 5 seconds exposure time per experiment. Fluorescence signal intensity along the first ~2 mm from the root tip was quantified using FUJI, and the background was averaged and subtracted for each image.

### Statistics and data processing

Student’s t-tests (two-tailed) were used to compare the statistical significance between pairs of samples. ANOVA and Turkey’s HSD were used to determine the statistical significance during multiple comparison using R (www.r-project.org). For CFU data, data were found to be normal after log transformation. All statistics were performed on the log-transformed data.

To compare the significance of the difference of gene expression between the parental line and *fer-8* (Extended data Fig. 9), we first ranked all the genes within the same sample according to their absolute expression values within a single genotype. For genes from each hormone pathway, we performed normal bootstrap on 5000 replicates, and calculated the mean sign of differential expression of all genes within a given hormone pathway (if expression of a gene is higher, same or lower in *fer-8* relative to the parental line, the sign would be 1, 0 and −1, respectively). We asked whether genes from a pathway are more likely to be upregulated in *fer-8* compared to the parental line relative to the mean sign of non-hormone-responsive genes. P values were calculated based on the normal distribution of mean signs from 5000 bootstrap dataset for each pathway.

Relative H2DCF values in Fig. 4e were calculated by normalizing raw values to the average of Col-0 from the respective independent experiment.

## Acknowledgments

We thank Fred Ausubel for critical reading of the manuscript and generously providing the *hsm* mutant collection, Alice Y. Cheung, Zhiyong Wang, Xin Li, Yuelin Zhang, and Geoffrey Wasteneys for kindly providing seed stocks, and Martin Hirst’s lab for assistance with sequencing.

## Author contributions

C.H.H. and Y.S. conceived of the project and designed experiments. Y.S. performed the majority of experiments and data analysis. X.C.Z. performed the previous *hsm* screen. A.W. analyzed the RNAseq and microbiome profiling data. D.T. conducted confocal microscopy imaging. Q.G. performed statistical analysis for the expression of hormone responsive genes and Pearson correlation assays. S.S., Y.L. and L.W. helped with gnotobioc plant assays. Experiments related to *mfec* were performed by R.S. with input from S.Y.H. C.H.H and Y.S. wrote the manuscript with input from all.

## Code availability

The code related to microbiome sequencing and RNAseq analysis are available on the Haney lab GitHub (https://github.com/haneylab/).

## Supplementary tables

Table S1. Expression of genes in the shoots and roots of *fer-8* and the parental line by RNAseq.

Table S2. Differentially expressed genes in *fer-8* relative to the parental line in shoots and roots.

Table S3. Enriched GO terms of highly-expressed genes in the shoots of *fer-8*.

Table S4. Expression of different hormone responsive genes.

## Extended data

**Extended Data Fig. 1.**
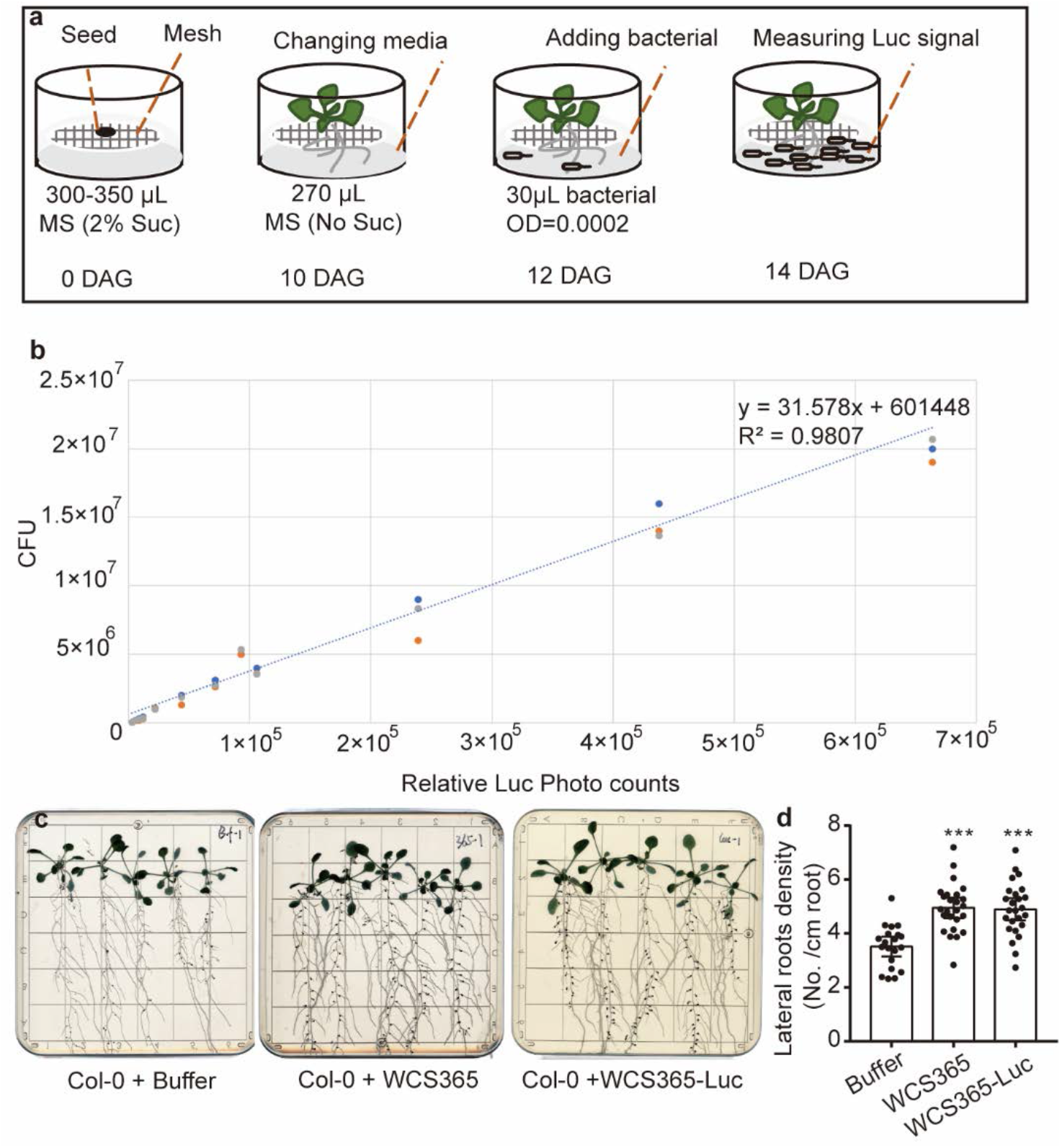
A gnotobiotic plant-rhizosphere microbe interaction system. **(a)** A diagram showing the 48-well plate based hydroponic plant-rhizosphere commensal interaction system. Seeds were grown on the Teflon mesh which separates shoots and roots after germination. 300-350 μL MS media with 2% sucrose was added to each well. Ten days after germination the media was replaced with 270 μL MS without sucrose. Two days later, 30 μL *P. fluorescens* WCS365-Luc (OD=0.0002) was added to the media, and rhizosphere bacteria levels were quantified after two days incubation using a plate reader. **(b)** WCS365-Luc CFU counts plotted against the relative Luc photo count signal intensities. For different dilutions, 100 μL WCS365-Luc (in 1× MS with 0.1% MES buffer) was transferred to a white opaque 96-well plate for quantifying Luc photo counts. A trend line and R^2^ were calculated in excel using the linear trendline option. Different colors of dots represent the three replicates of CFU plating results. **(c)** Insertion of the lux operon into WCS365 did not affect its ability to increase lateral root density. Images show plants treated with buffer, WCS365 or WCS365-Luc. **(d)** Quantification of lateral root density by normalizing to the number of lateral roots per cm of root. n=20, 25 and 25 from left to right. Mean ± SEM is shown, *** p<0.001 by Student’s t test compared to the buffer treated group.

**Extended Data Fig. 2.**
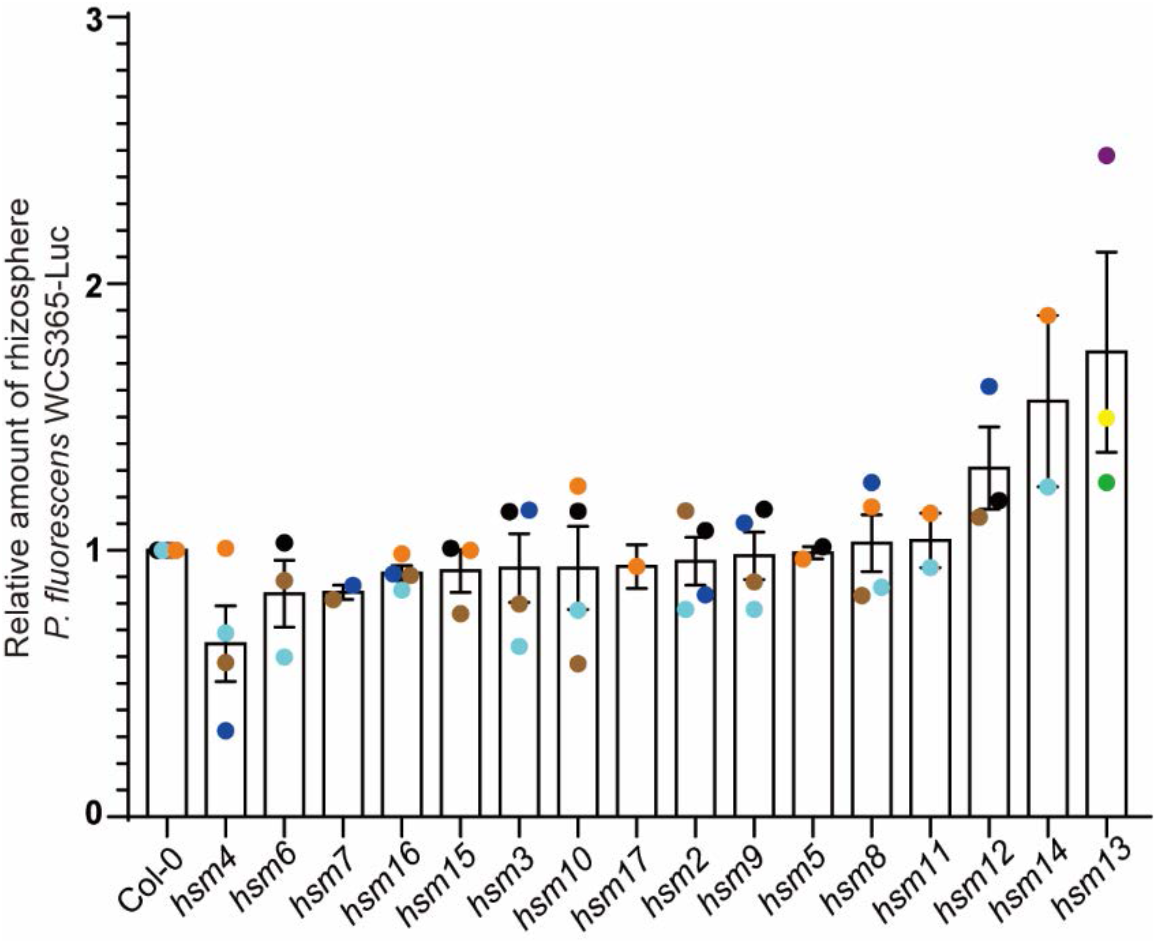
Relative amount (normalized to Col-0 for each experiment) of *P. fluorescens* WCS365-Luc in the rhizosphere of *hsm* mutants. Each dot represents the relative fold change of an *hsm* mutant relative to Col-0 from a single experiment; dots of the same color were performed as part of the same experimental replicate.

**Extended Data Fig. 3.**
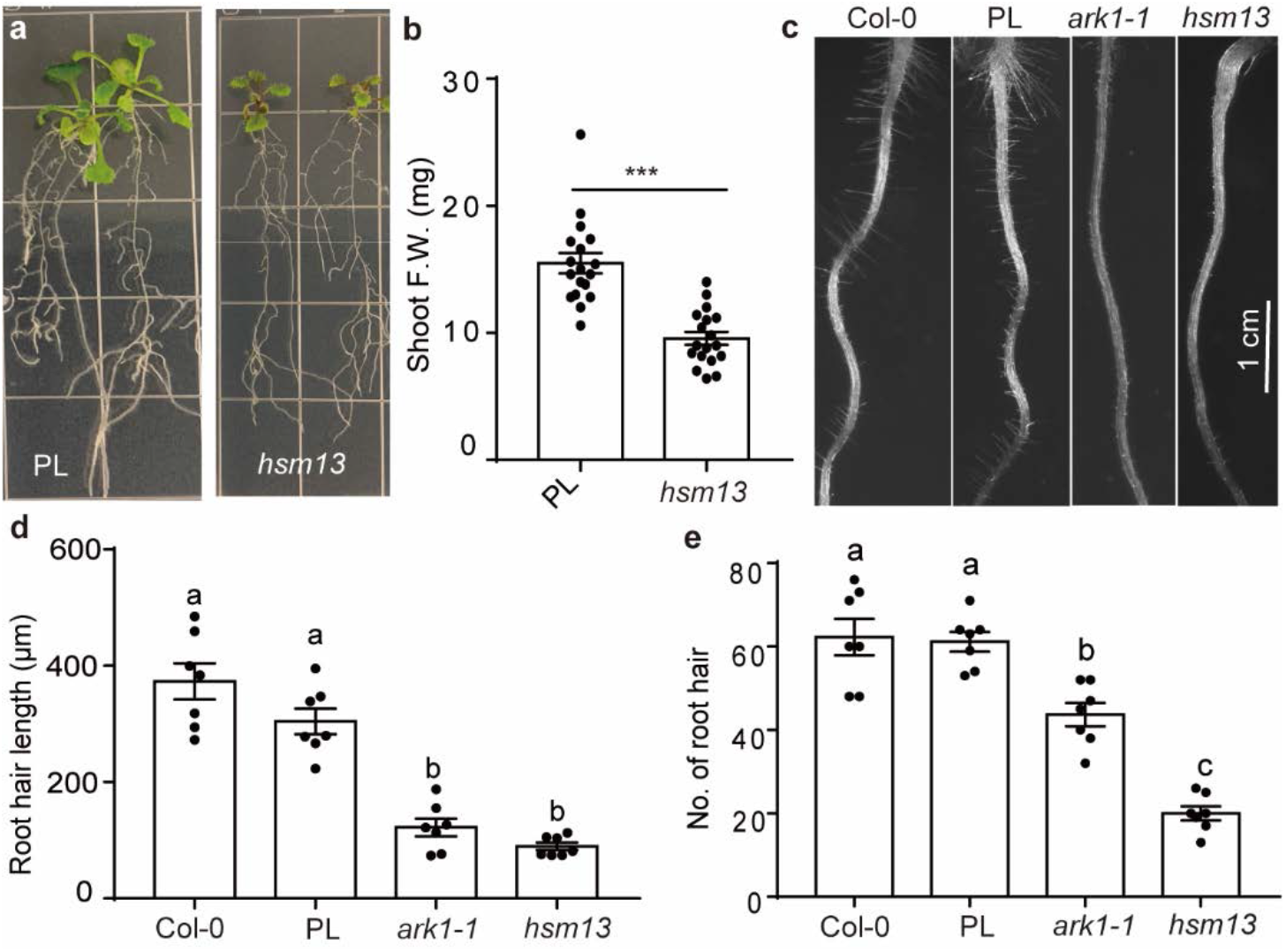
Morphology changes in *hsm13*. (**a-b**) *hsm13* is stunted when grown on axenic plates (½× MS agar) for 15 days. (**b**) Quantification of shoot fresh weight (F.W.) of *hsm13* and the parental line (PL) grown on plates. Student’s T-test was used to determine significance, and *** denotes p < 0.001. (**c**) *hsm13* shows a root hair developmental defect similar to the previously described *ark1-1* mutant. (**d-e**) Quantification of root hair length (**d**) and number of root hairs (**e**) from 6-day-old seedlings (0.5-4.5 mm region below the root-hypocotyl junction, n=7). Different letters indicate p < 0.05 by ANOVA and Turkey’s HSD test.

**Extended Data Fig. 4.**
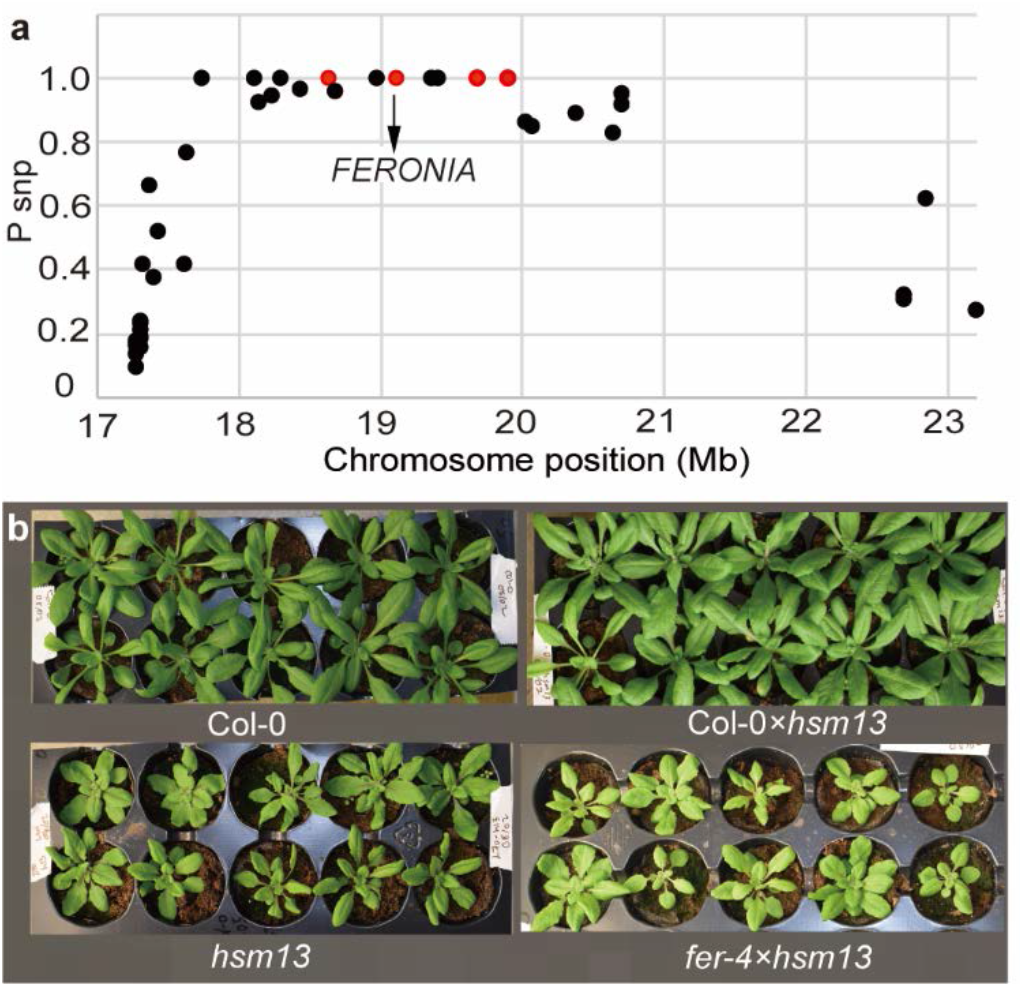
Mapping of *hsm13* based on the stunting phenotype. (**a**) Dot plot of SNP frequency on chromosome 3 in the *hsm13*-like F3s of *hsm13* × Col-0. Red dots represent non-synonymous mutations with P_SNP_=1 [P_SNP_=Relative reads abundance of mutant SNP/ (relative reads abundance of mutant SNP+ wildtype SNP)]. **(b)** F1 crosses between *fer-4* × *hsm13* are stunted while Col-0 × *hsm13* F1s had wildtype-like morphology.

**Extended Data Fig. 5.**
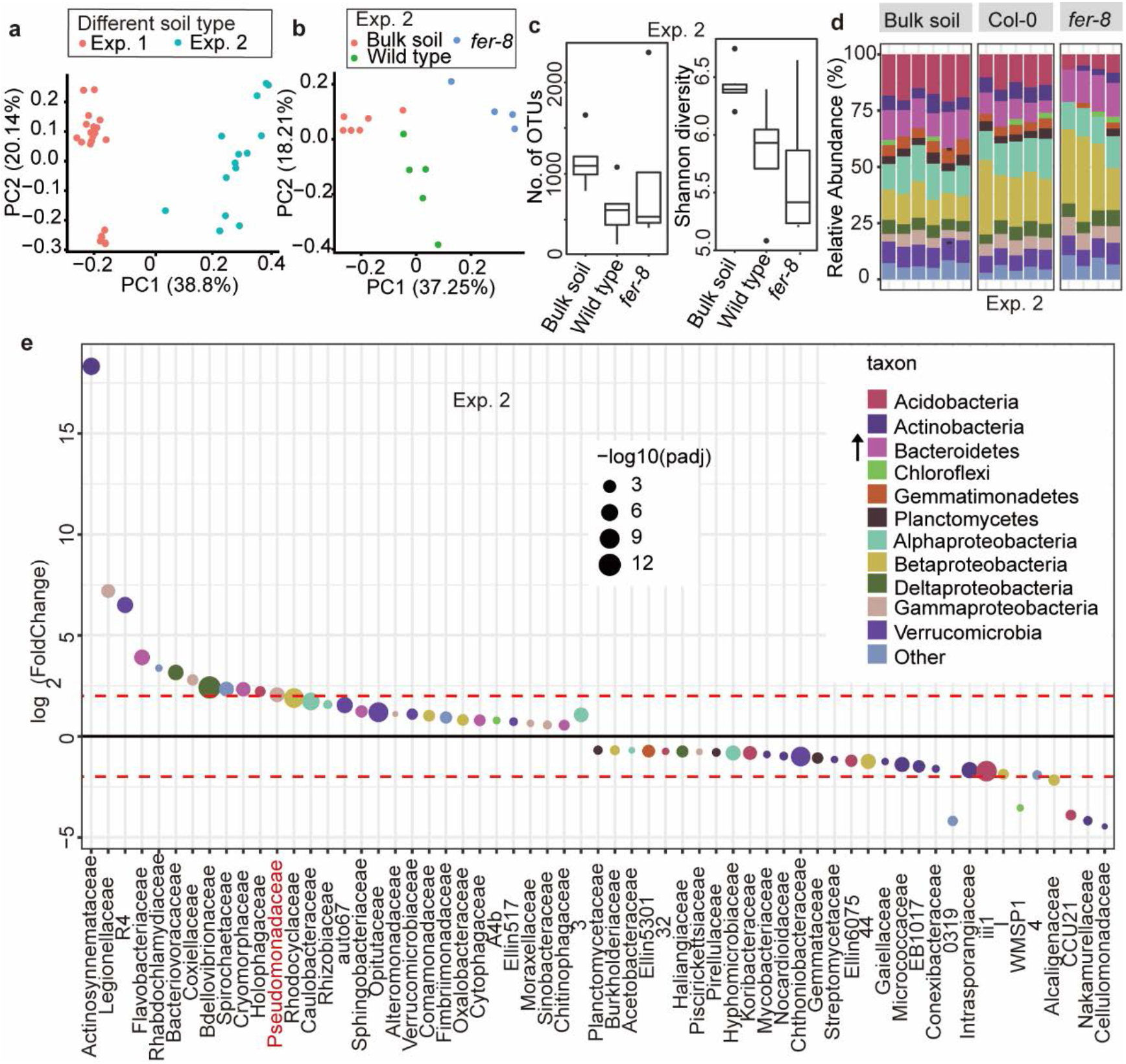
Pseudomonadaceae are robustly enriched in the rhizosphere of *fer-8* grown in soil from experiment 2. (**a**) All samples (bulk soil and plant rhizosphere) are from two independent experiments. Natural soil for Exp. 1 and 2 was harvest from the same site over two consecutive years. (**b**) Principal coordinate analysis (PCoA) shows *fer-8* has a distinct microbiome composition from Col-0 in Exp. 2. (**c**) Alpha diversity indexes in the bulk soil and rhizosphere samples (Exp. 2). (**d**) Relative abundance of bacterial phyla (or Proteobacterial classes) in bulk soil and different rhizosphere samples. The arrow shows the significantly changed taxon in the rhizosphere of *fer-8* relative to the parental line. **(e)** Significantly differentially abundant families between *fer-8* and wildtype rhizosphere. Colors show the taxon information for each family, and the dot size indicates the −log_10_ transformed adjusted p value; p<0.1 was used as a cutoff, red dash line indicates a 4-fold change [log2(fold change) =2].

**Extended Data Fig. 6.**
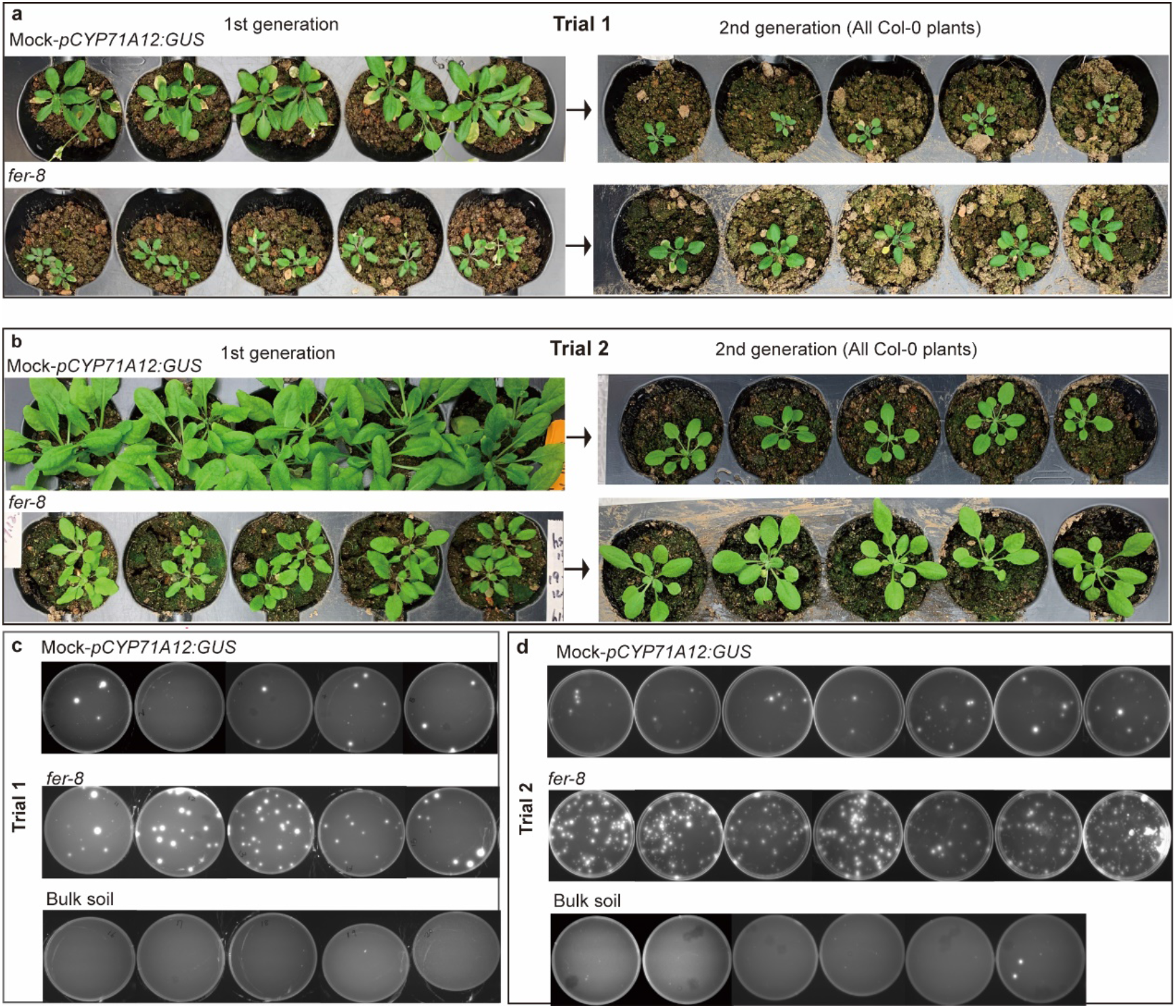
Phenotypes of plants from microbiome transplant experiments. **(a-b)** Images of 5-week old parental line (PL) and *fer-8* plants grown in natural soil (left panel), and images of 3-week old 2nd generation wildtype plants grown in the soil with previous cultivation of *fer-8* or PL (right panel). **(c-d)** Plating of rhizosphere samples from bulk soil, G1-*fer-8* and G1-PL plants. 100 μl of 0.00025 g/ml serially diluted sample was plated on King’s B media and imaged under UV light.

**Extended Data Fig. 7.**
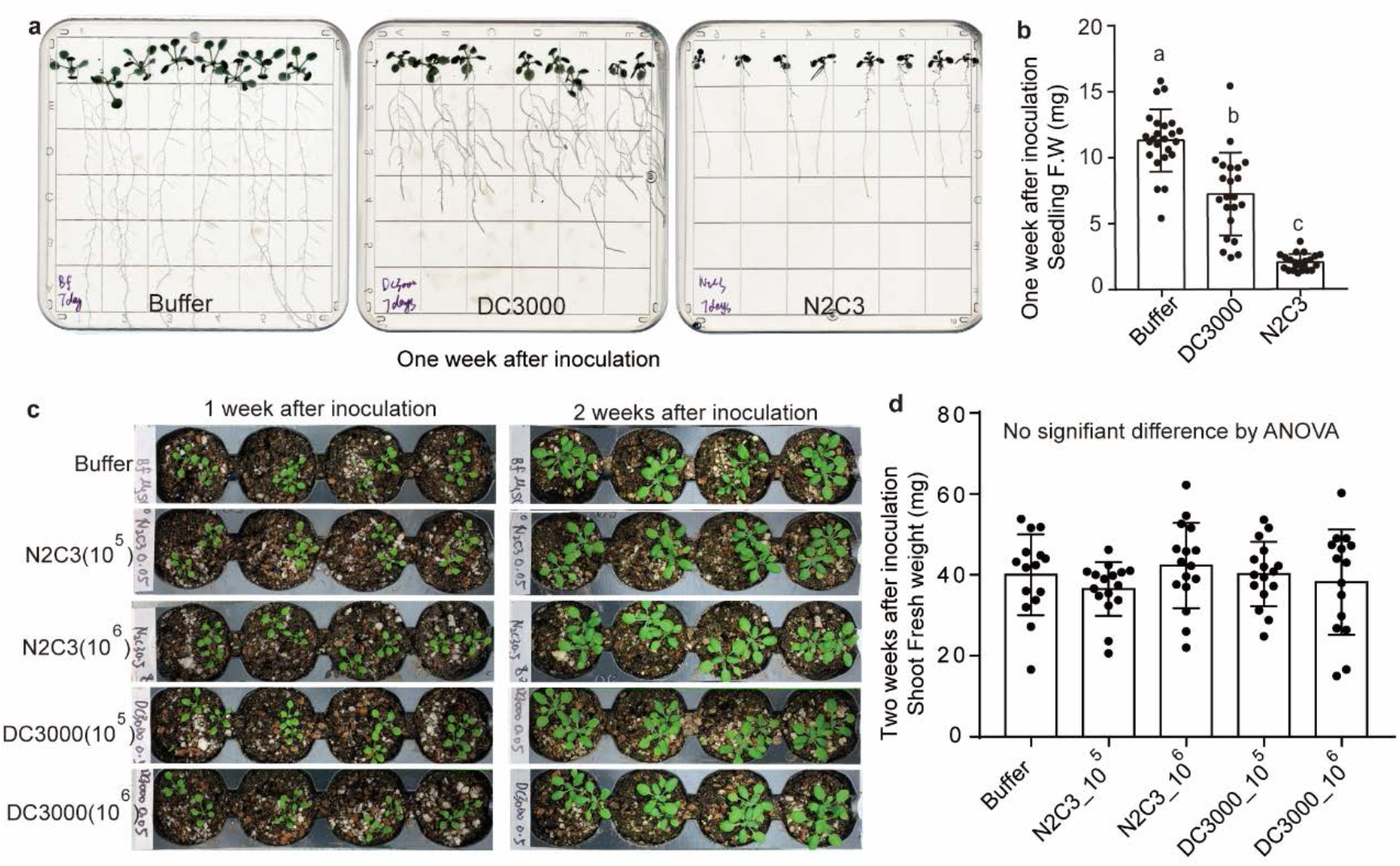
*Pseudomonas* pathogens do not efficiently cause disease in soil. (**a**) Under axenic conditions, *Pst* DC3000 and *Pseudomonas* sp. N2C3 cause stunting one week after inoculation. Seeds were grown on ½× MS without sucrose for 6 days before inoculation. (**b**) Quantification of seedling fresh weight (F.W.) one week after inoculation on plates. (b and d) Mean ± SEM is shown, different letters indicate p < 0.05 by ANOVA and Turkey’s HSD test. (**c**) Phenotypes of Col-0 grown in natural soil 1 and 2 weeks after rhizosphere inoculation with *Pst* DC3000 or *Pseudomonas* sp. N2C3. For both pathogens, 1 mL OD_600_=0.5 or 0.05 inoculum was added to reach a final concentration of 10^5^ or 10^6^ CFU/gram soil. (**d**) Quantification of shoot fresh weight 2 weeks after inoculation.

**Extended Data Fig. 8.**
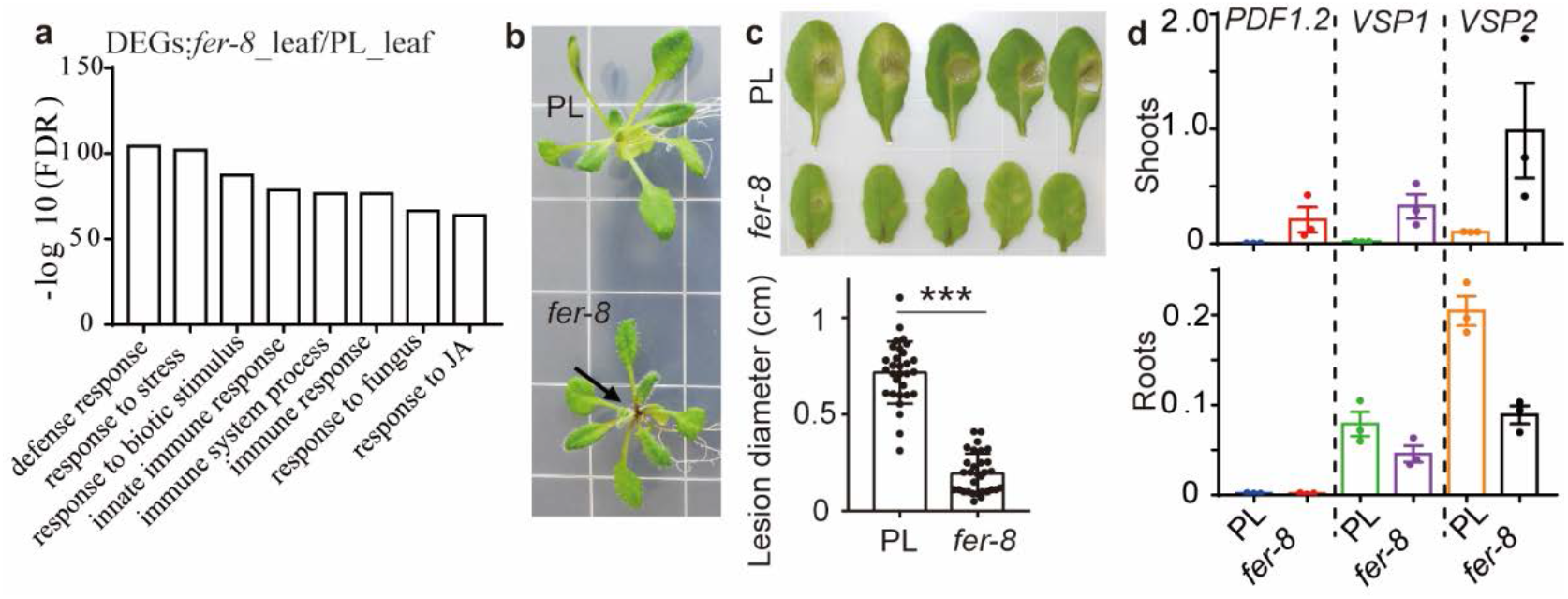
JA signaling is elevated in the shoots of *fer-8*. (**a**) Selected Gene Ontology categories from top 20 significantly enriched GO terms (based on the −log transformed FDR values). GO enrichment analysis was performed with AgriGO. (**b**) Anthocyanin accumulation was observed in the petioles of *fer-8* but not the parental line (PL); representative 2-week old plants grown on plates are shown. (**c**) *fer-8* is resistant to the necrotrophic pathogen *Botrytis cinerea*. The image shows the lesion size 3 days after inoculation. n = 30 and 31 for the PL and *fer-8*, respectively. Data from 2 independent experiments were used for analysis. *** p <0.001 by Student’s t-test. (**d**) JA responsive genes were highly expressed in the shoots of *fer-8* but not in the roots. qRT-PCR data from three biological replicates. Gene expression values were normalized to the expression of an internal control gene *EIF4A*. Mean ± SEM is shown.

**Extended Data Fig. 9.**
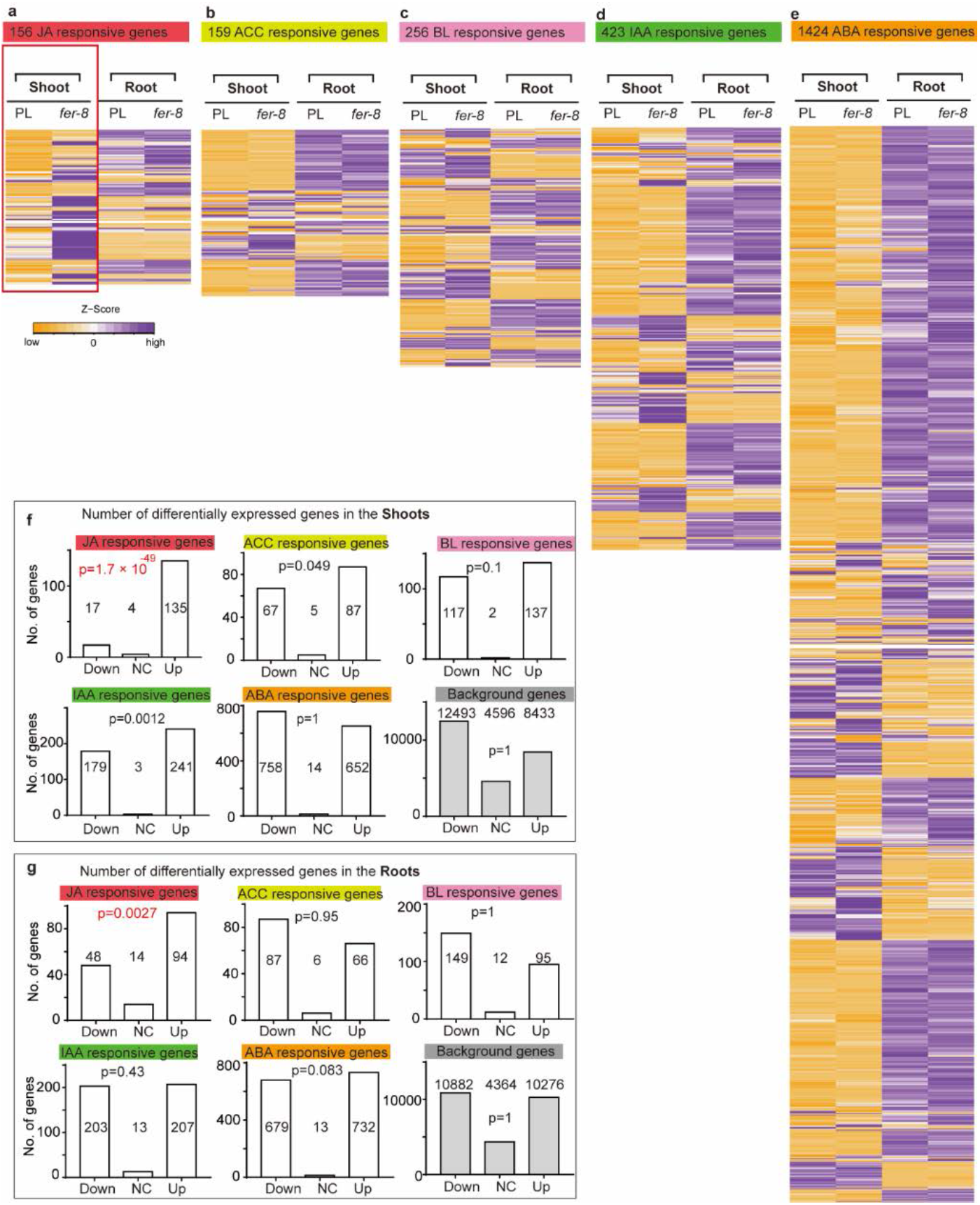
The effect of the *fer*-8 mutation on hormone signaling is shoot specific and most significantly related to JA signaling. **(a-e)** The heatmaps show the normalized expression [Z score of TPMs (Transcripts Per Million), Z score = (experimental score – mean)/standard deviation] of hormone-responsive genes in the shoots and roots of *fer-8* and the parental line (PL). JA: Jasmonic acid; ACC: 1-aminocyclopropane-1-carboxylate, an ethylene biosynthetic precursor; BL: brassinolide; IAA: indole-3-acetic acid; ABA: abscisic acid. Each column shows the average values from different replicates within each genotype and tissue. Seedlings were grown under axenic conditions (½× MS-Agar plates). Gene list and expression values for different groups are listed in the Supplementary table S4. (**f-g**) Comparing the rank difference of genes from different hormone responsive pathways and the genome background (non-hormone responsive genes) in shoots and roots. Up, Down and NC means the number of genes with a higher, lower or same ranked in *fer-8* relative to PL, respectively. The p-values were calculated by performing normal bootstrap (5000 replicates) on the mean sign (sign=1, 0 and −1 if a gene is higher, same or lower ranked in *fer-8*) of the rank difference within each pathway (Methods).

**Extended Data Fig. 10.**
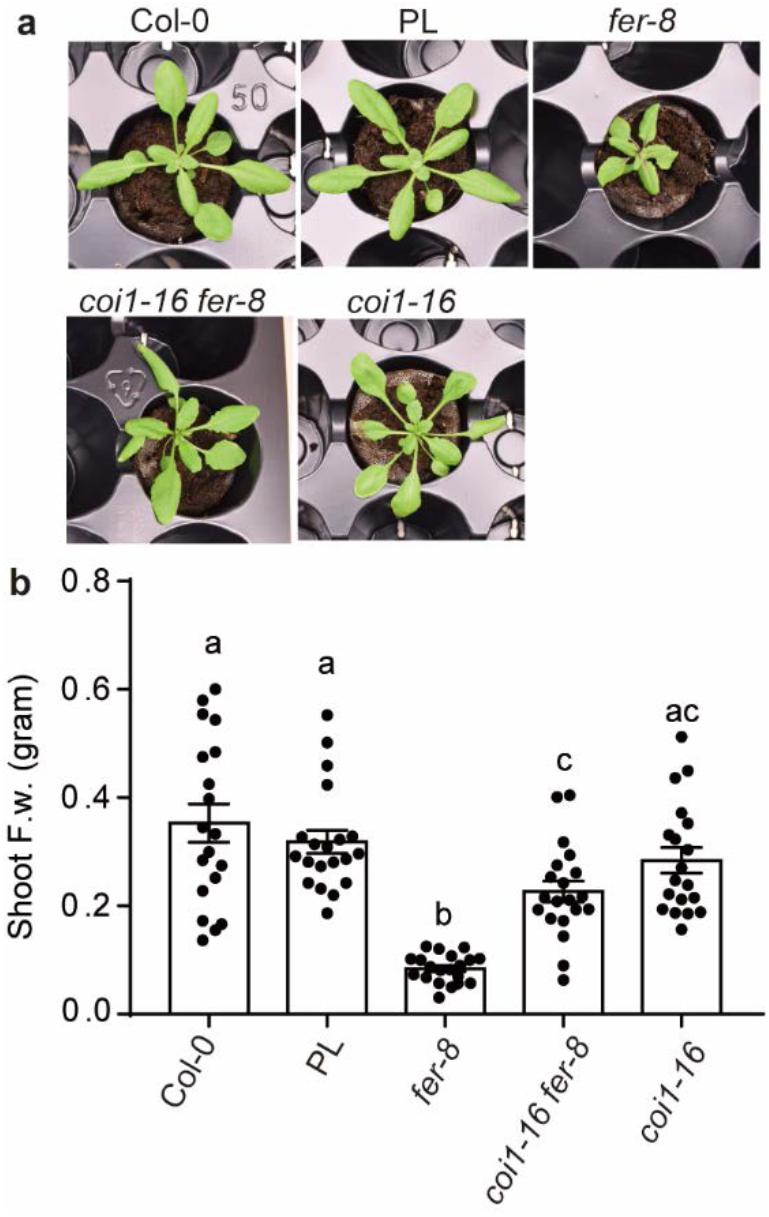
*fer-8 coi1-16* double mutant suppressed the stunting phenotype of *fer-8*. (**a**) Growth phenotypes of Col-0, the parental line (PL), *fer-8, coi1-16 fer-8* and *coi1-16*. Images were taken three weeks after germination; (**b**) Quantification of shoot fresh weight (F.w.) of different genotypes at three weeks after germination. Mean ± SEM, different letters indicate p < 0.05 by ANOVA and Turkey’s HSD test.

**Extended Data Fig. 11.**
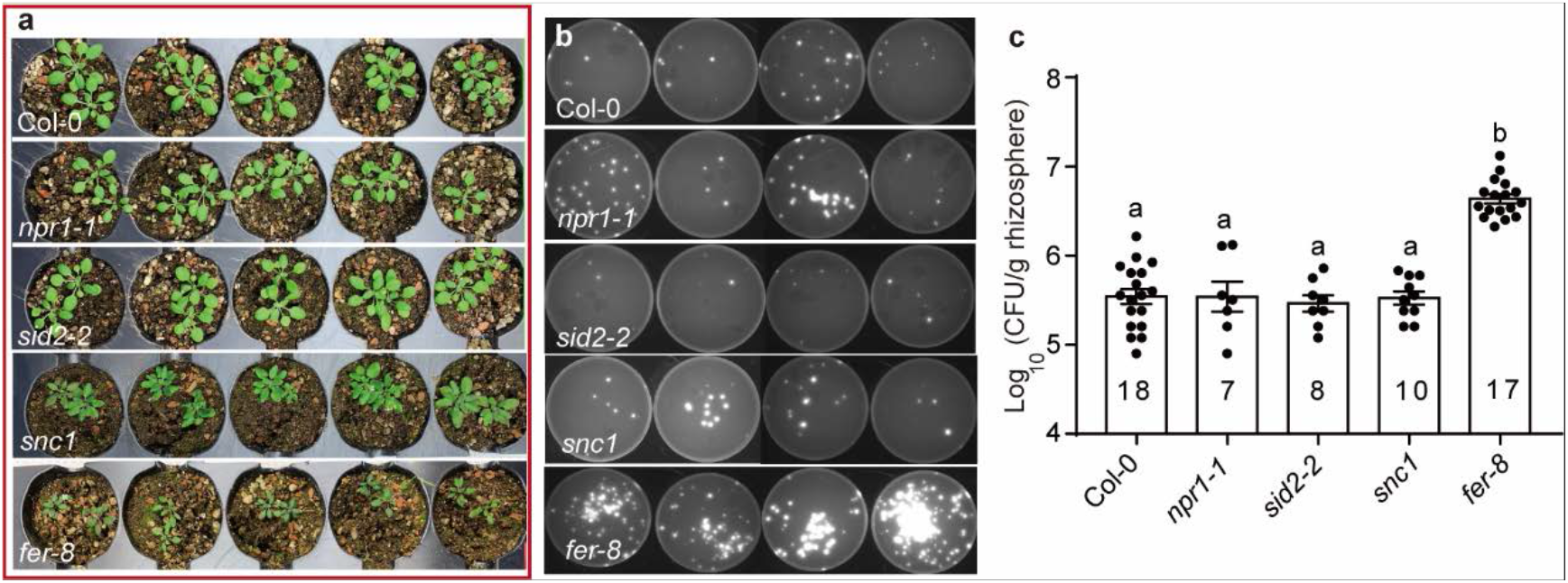
Salicylic acid (SA) mutants do not affect levels of rhizosphere fluorescent Pseudomonads. (**a**) Phenotypes of Col-0, SA signaling and biosynthesis mutants *npr1-1, sid2-2, snc1*, and *fer-8* grown in natural soil. (**b**) Representative plates of rhizosphere samples from genotypes in (a) are shown. (**c**) Quantification of rhizosphere fluorescent Pseudomonads. Mean ± SEM, different letters indicate p < 0.05 by ANOVA and Turkey’s HSD test. Numbers represent the biological replicates from 2-4 independent experiments.

**Extended Data Fig. 12.**
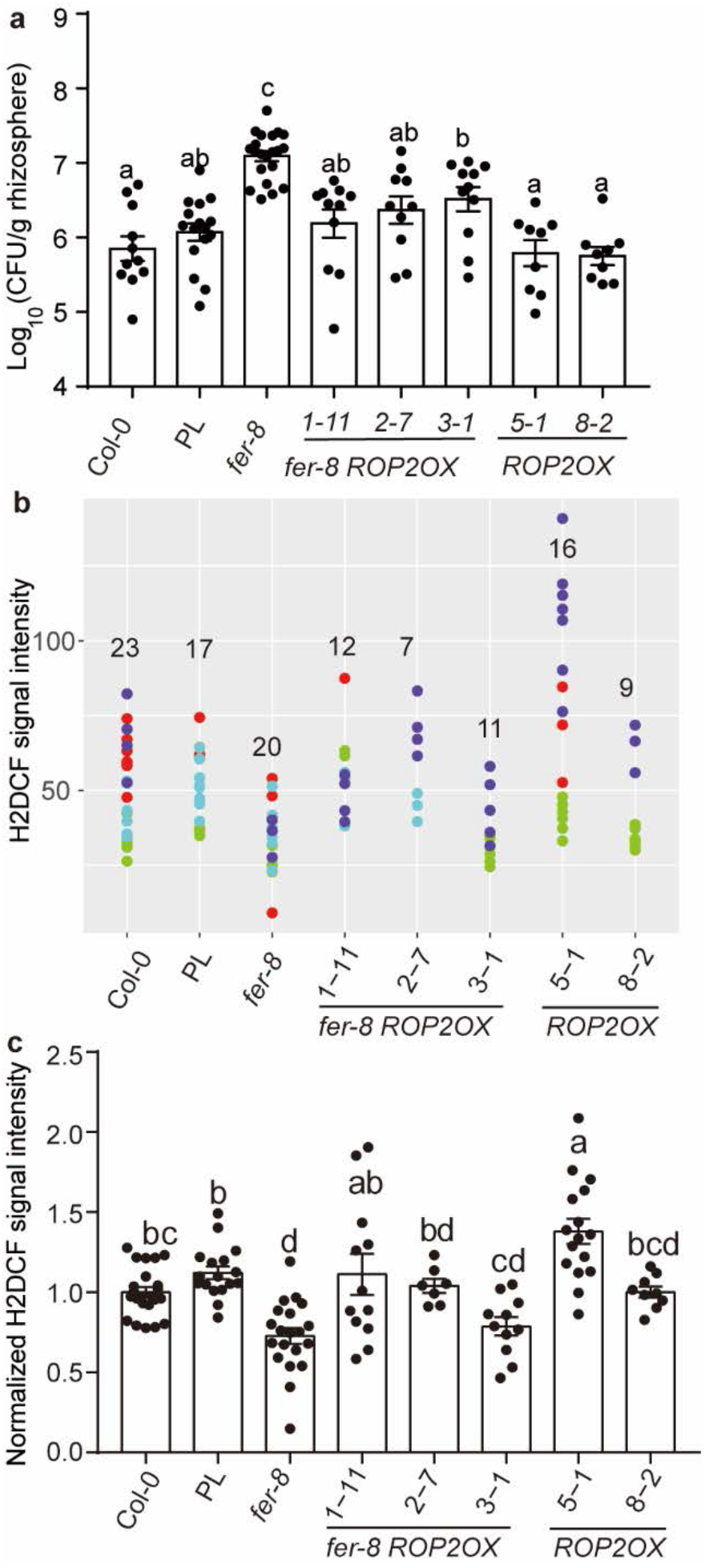
H2DCF staining of ROS levels in roots. (**a**) Overexpression of *ROP2* (a positive regulator of NADPH oxidase) in *fer-8* decreases rhizosphere levels of fluorescent Pseudomonads. n=11, 16, 21, 11, 10, 11, 9 and 9 from left to right (3-5 independent experiments). (**b**) Signal intensity values of root H2DCF staining results from 2-4 independent experiments for Col-0, *fer-8* and *fer-8 ROP2OX* (line 1-11, 5-1 and 3-1), and *ROP2OX* (line8*-2* and 5-1). Numbers on the graph denote the number of measured plants, and dots of the same color were performed as part of the same experimental replicate. (**c**) Data from different independent experiments were normalized to the average values of the Col-0 control from the same experiment. Mean ± SEM, different letters indicate p < 0.05 by ANOVA and Turkey’s HSD test.

## Main references

1 Chen, Tao, et al. “A plant genetic network for preventing dysbiosis in the phyllosphere.” Nature 580, 653–657. (2020)

2 Levy, M., Kolodziejczyk, A. A., Thaiss, C. A. & Elinav, E. Dysbiosis and the immune system. Nat Rev Immunol 17, 219–232, (2017).

3 de Vries, F. T., Griffiths, R. I., Knight, C. G., Nicolitch, O. & Williams, A. Harnessing rhizosphere microbiomes for drought-resilient crop production. Science 368, 270–274, (2020).

4 Berendsen, R. L., Pieterse, C. M. & Bakker, P. A. The rhizosphere microbiome and plant health. Trends in plant science 17, 478–486 (2012).

5 Duan, Q., Kita, D., Li, C., Cheung, A. Y. & Wu, H. M. FERONIA receptor-like kinase regulates RHO GTPase signaling of root hair development. PNAS 107, 17821–17826, (2010).

6 Rosenberg, E. & Zilber-Rosenberg, I. Microbes Drive Evolution of Animals and Plants: the Hologenome Concept. MBio 7, e01395, (2016).

7 Lugtenberg, B. & Kamilova, F. Plant-growth-promoting rhizobacteria. Annual review of microbiology 63, 541–556, (2009).

8 Zhang, J. et al. *NRT1.1B* is associated with root microbiota composition and nitrogen use in field-grown rice. Nat Biotechnol, (2019).

9 Kwak, M. J. et al. Rhizosphere microbiome structure alters to enable wilt resistance in tomato. Nat Biotechnol, (2018).

10 Dimkpa, C., Weinand, T. & Asch, F. Plant-rhizobacteria interactions alleviate abiotic stress conditions. Plant, cell & environment 32, 1682–1694, (2009).

11 Abdelaziz, M. E. et al. Piriformospora indica alters Na+/K+ homeostasis, antioxidant enzymes and LeNHX1 expression of greenhouse tomato grown under salt stress. Scientia Horticulturae 256, (2019).

12 Berendsen, R. L. et al. Disease-induced assemblage of a plant-beneficial bacterial consortium. The ISME journal 12, 1496–1507, (2018).

13 Mendes, R. et al. Deciphering the rhizosphere microbiome for disease-suppressive bacteria. Science 332, 1097–1100 (2011).

14 Vurukonda, S. S., Vardharajula, S., Shrivastava, M. & Sk, Z. A. Enhancement of drought stress tolerance in crops by plant growth promoting rhizobacteria. Microbiol Res 184, 13–24, (2016).

15 Haas, D. & Defago, G. Biological control of soil-borne pathogens by fluorescent Pseudomonads. Nat Rev Microbiol 3, 307–319, (2005).

16 Bakker, P. A., Pieterse, C. M. & van Loon, L. C. Induced Systemic Resistance by Fluorescent *Pseudomonas* spp. Phytopathology 97, 239–243, (2007).

17 Weller, D. M., Howie, W. J. & Cook, R. J. Relationship between *in vitro* inhibition of *Gaeumannomyces graminis* var. *tritici* and suppression of take-all of wheat by fluorescent Pseudomonads. Phytopathology 78, 1094–1100 (1988).

18 Mazurier, S., Corberand, T., Lemanceau, P. & Raaijmakers, J. M. Phenazine antibiotics produced by fluorescent Pseudomonads contribute to natural soil suppressiveness to *Fusarium* wilt. The ISME journal 3, 977 (2009).

19 Stutz, E., Défago, G. & Kern, H. Naturally Occurring Fluorescent Pseudomonads Involved in Suppression of black root rot of tobacco. Phytopathology 76, 181–185 (1986).

20 Shipton, P. Monoculture and soilborne plant pathogens. Annual review of phytopathology 15, 387–407 (1977).

21 Simon, A. & Sivasithamparam, K. Pathogen-suppression: a case study in biological suppression of *Gaeumannomyces graminis* var. *tritici* in soil. Soil Biology and Biochemistry 21, 331–337 (1989).

22 Cook, R. J. The influence of rotation crops on take-all decline phenomenon. Phytopathology 71, 189–192 (1981).

23 Haney, C. H., Samuel, B. S., Bush, J. & Ausubel, F. M. Associations with rhizosphere bacteria can confer an adaptive advantage to plants. Nature plants 1, (2015).

24 Zhang, X. C., Millet, Y. A., Cheng, Z., Bush, J. & Ausubel, F. M. Jasmonate signalling in *Arabidopsis* involves SGT1b-HSP70-HSP90 chaperone complexes. Nature plants 1, (2015).

25 Johnsen, K. & Nielsen, P. Diversity of *Pseudomonas* strains isolated with King’s B and Gould’s S1 agar determined by repetitive extragenic palindromic-polymerase chain reaction, 16S rDNA sequencing and Fourier transform infrared spectroscopy characterisation. FEMS microbiology letters 173, 155–162, (1999).

26 Noguchi, T. et al. Brassinosteroid-insensitive dwarf mutants of *Arabidopsis* accumulate brassinosteroids. Plant Physiol 121, 743–752, (1999).

27 Eng, R. C. & Wasteneys, G. O. The microtubule plus-end tracking protein ARMADILLO-REPEAT KINESIN1 promotes microtubule catastrophe in *Arabidopsis*. Plant Cell 26, 3372–3386, (2014).

28 Guo, H. et al. FERONIA Receptor Kinase Contributes to Plant Immunity by Suppressing Jasmonic Acid Signaling in *Arabidopsis thaliana*. Curr Biol 28, 3316–3324 e3316, (2018).

29 Haruta, M., Sabat, G., Stecker, K., Minkoff, B. B. & Sussman, M. R. A peptide hormone and its receptor protein kinase regulate plant cell expansion. Science 343, 408–411, (2014).

30 Kim, B. R. et al. Deciphering Diversity Indices for a Better Understanding of Microbial Communities. J Microbiol Biotechnol 27, 2089–2093, (2017).

31 Melnyk, R. A., Hossain, S. S. & Haney, C. H. Convergent gain and loss of genomic islands drive lifestyle changes in plant-associated *Pseudomonas*. The ISME journal 13, 1575–1588, (2019).

32 Haney, C. H. et al. Rhizosphere-associated *Pseudomonas* induce systemic resistance to herbivores at the cost of susceptibility to bacterial pathogens. Mol Ecol 27, 1833–1847, (2018).

33 Carrion, V. J. et al. Pathogen-induced activation of disease-suppressive functions in the endophytic root microbiome. Science 366, 606–612, (2019).

34 Raaijmakers, J. M. et al. Dose-response relationships in biological control of *Fusarium* wilt of radish by *Pseudomonas* spp. Phytopathology 85, 1075–1080 (1995).

35 Ellis, C. & Turner, J. G. A conditionally fertile *coi1* allele indicates cross-talk between plant hormone signalling pathways in *Arabidopsis thaliana* seeds and young seedlings. Planta 215, 549–556, (2002).

36 Spoel, S. H. et al. NPR1 modulates cross-talk between salicylate- and jasmonate-dependent defense pathways through a novel function in the cytosol. Plant Cell 15, 760–770, (2003).

37 Cao, H., Bowling, S. A., Gordon, A. S. & Dong, X. Characterization of an *Arabidopsis* Mutant That Is Nonresponsive to Inducers of Systemic Acquired Resistance. Plant Cell 6, 1583–1592, (1994).

38 Dewdney, J. et al. Three unique mutants of *Arabidopsis* identify *eds* loci required for limiting growth of a biotrophic fungal pathogen. The Plant journal 24, 205–218, (2000).

39 Zhang, Y., Goritschnig, S., Dong, X. & Li, X. A gain-of-function mutation in a plant disease resistance gene leads to constitutive activation of downstream signal transduction pathways in *suppressor of npr1-1, constitutive 1*. Plant Cell 15, 2636–2646, (2003).

40 Stegmann, M. et al. The receptor kinase FER is a RALF-regulated scaffold controlling plant immune signaling. Science 355, 287–289, (2017).

41 Miller, G. et al. The plant NADPH oxidase RBOHD mediates rapid systemic signaling in response to diverse stimuli. Science signaling 2, ra45, (2009).

42 Zhou, F. et al. Co-incidence of Damage and Microbial Patterns Controls Localized Immune Responses in Roots. Cell 180, 440–453 e418, (2020).

43 Yu, K. et al. Rhizosphere-Associated *Pseudomonas* Suppress Local Root Immune Responses by Gluconic Acid-Mediated Lowering of Environmental pH. Curr Biol 29, 3913–3920 e3914, (2019).

44 Masachis, S. et al. A fungal pathogen secretes plant alkalinizing peptides to increase infection. Nat Microbiol 1, 16043, (2016).

45 Zhang, Xin, et al. Nematode-encoded RALF peptide mimics facilitate parasitism of plants through the FERONIA receptor kinase. Molecular Plant 13, 1434–1454, (2020).

46 Hussain, M. et al. Bacterial community assemblages in the rhizosphere soil, root endosphere and cyst of soybean cyst nematode-suppressive soil challenged with nematodes. FEMS microbiology ecology 94, (2018).

47 Zhang, X., Yang, Z., Wu, D. & Yu, F. RALF–FERONIA Signaling: Linking Plant Immune Response with Cell Growth. Plant Communications 1, (2020).

48 Campbell, L. & Turner, S. R. A Comprehensive Analysis of RALF Proteins in Green Plants Suggests There Are Two Distinct Functional Groups. Frontiers in Plant Science 8, (2017).

## Methods references

49 Gómez-Gómez, L. & Boller, T. FLS2: an LRR receptor-like kinase involved in the perception of the bacterial elicitor flagellin in *Arabidopsis*. Mol Cell 5, 1003–1011, (2000).

50 Zipfel, C. et al. Perception of the bacterial PAMP EF-Tu by the receptor EFR restricts *Agrobacterium-mediated* transformation. Cell 125, 749–760, (2006).

51 Schwessinger, B. et al. Phosphorylation-dependent differential regulation of plant growth, cell death, and innate immunity by the regulatory receptor-like kinase BAK1. PLoS genetics 7, e1002046 (2011).

52 Lane, M. C., Alteri, C. J., Smith, S. N. & Mobley, H. L. Expression of flagella is coincident with uropathogenic *Escherichia coli* ascension to the upper urinary tract. PNAS 104, 16669–16674 (2007).

53 James, G. V. et al. User guide for mapping-by-sequencing in *Arabidopsis*. Genome biology 14, R61, (2013).

54 Li, H. & Durbin, R. Fast and accurate short read alignment with Burrows-Wheeler transform. Bioinformatics 25, 1754–1760, (2009).

55 Callahan, B. J. et al. DADA2: High-resolution sample inference from Illumina amplicon data. Nature methods 13, 581–583, (2016).

56 Love, M. I., Huber, W. & Anders, S. Moderated estimation of fold change and dispersion for RNA-seq data with DESeq2. Genome biology 15, 550, (2014).

57 Ding, Y. et al. Opposite Roles of Salicylic Acid Receptors NPR1 and NPR3/NPR4 in Transcriptional Regulation of Plant Immunity. Cell 173, 1454–1467.e1415, (2018).

58 Geels, F. & Schippers, B. Selection of antagonistic fluorescent *Pseudomonas* spp. and their root colonization and persistence following treatment of seed potatoes. Journal of Phytopathology 108, 193–206 (1983).

59 Lamers, J., Schippers, B. & Geels, F. Soil-borne diseases of wheat in the Netherlands and results of seed bacterization with pseudomonads against *Gaeumannomyces graminis* var. *tritici*, associated with disease resistance. Cereal breeding related to integrated cereal production, 134–139 (1988).

60 Holloway, B. W. Genetic recombination in *Pseudomonas aeruginosa*. Journal of general microbiology 13, 572–581, (1955).

61 Earl, A. M., Losick, R. & Kolter, R. *Bacillus subtilis* genome diversity. Journal of bacteriology 189, 1163–1170, (2007).

62 Bedard, D. L. et al. Rapid assay for screening and characterizing microorganisms for the ability to degrade polychlorinated biphenyls. Applied and environmental microbiology 51, 761–768 (1986).

63 Baldani, J., Baldani, V., Seldin, L. & Döbereiner, J. Characterization of *Herbaspirillum seropedicae* gen. nov., sp. nov., a root-associated nitrogen-fixing bacterium. International Journal of Systematic and Evolutionary Microbiology 36, 86–93 (1986).

64 Sessitsch, A., et al. *Burkholderia phytofirmans* sp. nov., a novel plant-associated bacterium with plant-beneficial properties. Int. J. Syst. Evol. Microbiol. 55, 1187–1192, (2005).

65 Cuppels, D. A. Generation and Characterization of *Tn5* Insertion Mutations in *Pseudomonas syringae* pv. *tomato*. Applied and environmental microbiology 51, 323–327, (1986).

66 Price, M. N. et al. Deep annotation of protein function across diverse bacteria from mutant phenotypes. BioRxiv, 072470 (2016).

67 Langmead, B., Trapnell, C., Pop, M. & Salzberg, S. L. Ultrafast and memory-efficient alignment of short DNA sequences to the human genome. Genome biology 10, R25, (2009).

68 Li, B. & Dewey, C. N. RSEM: accurate transcript quantification from RNA-Seq data with or without a reference genome. BMC bioinformatics 12, 323, (2011).

69 Tian, T. et al. agriGO v2.0: a GO analysis toolkit for the agricultural community, 2017 update. Nucleic acids research 45, W122–w129, (2017).

70 Hickman, R. et al. Architecture and Dynamics of the Jasmonic Acid Gene Regulatory Network. Plant Cell 29, 2086–2105, (2017).

71 Nemhauser, J. L., Hong, F. & Chory, J. Different plant hormones regulate similar processes through largely nonoverlapping transcriptional responses. Cell 126, 467–475, (2006).

72 Genenncher, B. et al. Nucleoporin-Regulated MAP Kinase Signaling in Immunity to a Necrotrophic Fungal Pathogen. Plant Physiol 172, 1293–1305, (2016).

73 Kirchhelle, C. & Moore, I. A simple chamber for long-term confocal imaging of root and hypocotyl development. JoVE (Journal of Visualized Experiments), e55331 (2017).

74 Schindelin, J. et al. Fiji: an open-source platform for biological-image analysis. Nature methods 9, 676–682, (2012).

